# LONP1 regulation of mitochondrial protein folding provides insight into beta cell failure in type 2 diabetes

**DOI:** 10.1101/2024.06.03.597215

**Authors:** Jin Li, Jie Zhu, Yamei Deng, Emma C. Reck, Emily M. Walker, Vaibhav Sidarala, Dre L. Hubers, Mabelle B. Pasmooij, Chun-Shik Shin, Khushdeep Bandesh, Eftyhmios Motakis, Siddhi Nargund, Romy Kursawe, Venkatesha Basrur, Alexey I. Nesvizhskii, Michael L. Stitzel, David C. Chan, Scott A. Soleimanpour

## Abstract

Proteotoxicity is a contributor to the development of type 2 diabetes (T2D), but it is unknown whether protein misfolding in T2D is generalized or has special features. Here, we report a robust accumulation of misfolded proteins within the mitochondria of human pancreatic islets in T2D and elucidate its impact on β cell viability. Surprisingly, quantitative proteomics studies of protein aggregates reveal that human islets from donors with T2D have a signature more closely resembling mitochondrial rather than ER protein misfolding. The matrix protease LonP1 and its chaperone partner mtHSP70 were among the proteins enriched in protein aggregates. Deletion of LONP1 in mice yields mitochondrial protein misfolding and reduced respiratory function, ultimately leading to β cell apoptosis and hyperglycemia. Intriguingly, LONP1 gain of function ameliorates mitochondrial protein misfolding and restores human β cell survival following glucolipotoxicity via a protease-independent effect requiring LONP1-mtHSP70 chaperone activity. Thus, LONP1 promotes β cell survival and prevents hyperglycemia by facilitating mitochondrial protein folding. These observations may open novel insights into the nature of impaired proteostasis on β cell loss in the pathogenesis of T2D that could be considered as future therapeutic targets.

## INTRODUCTION

Optimal protein homeostasis, also known as proteostasis, is vital to combating aging-related diseases, including cancer, cardiovascular disorders, neurodegenerative diseases, and type 2 diabetes (T2D).^1, 2^ Impairments in proteostasis include a decline in the coordinated balance of protein synthesis, proper folding, structural maintenance, and turnover.^3, 4^ Emerging evidence demonstrates that T2D is a protein misfolding disease.^5^ Among the best examples of protein misfolding in T2D occur in pancreatic β-cells, where misfolding of proinsulin and islet amyloid polypeptide (IAPP), a hormone co-secreted with insulin, lead to β-cell dysfunction.^6–11^ Proinsulin misfolding precipitates endoplasmic reticulum (ER) stress, while aggregates of IAPP lead to a cascade of defects culminating in β-cell apoptosis.^6–11^ Accordingly, ER protein misfolding and stress is often considered a major mediator of β-cell apoptosis in T2D.^12, 13^ Beyond proinsulin and IAPP, however, the extent, location, and specific impact of protein misfolding in T2D is unclear.

A parallel and related feature to proteostasis of importance to aging-related diseases and T2D is the development of mitochondrial dysfunction. Mitochondria are crucial for several vital functions in pancreatic β-cells including support of fuel-stimulated insulin release and maintenance of β-cell mass and survival.^14–17^ Indeed, β cells from islet donors with T2D develop dilated mitochondrial ultrastructure with dysmorphic cristae as well as bioenergetic defects.^18, 19^ The observation of increases in reactive oxygen species (ROS) and β cell oxidative damage in individuals with T2D may also be closely related to mitochondrial damage, as mitochondria are a major source of ROS production.^20^ Further, recent work has identified that β cell mitochondrial gene expression and oxidative phosphorylation defects precede the development of T2D.^21^ Human genetic studies also support associations between mitochondria and T2D.^22–28^ However, a mechanistic link between impairments in β cell proteostasis and mitochondrial health in T2D has not been examined.

The majority of the mitochondrial proteome is encoded by the nuclear genome and synthesized on cytosolic ribosomes. Optimal mitochondrial proteostasis depends on proper import and folding of unfolded mitochondrial precursors into their functional structures as well as safeguard mechanisms to respond to protein misfolding or stress. Within the mitochondrial matrix, chaperone complexes comprised of mitochondrial (mt)HSP70 (also known as HSPA9/GRP75/mortalin), DNAJA3, and GRPEL1/2 as well as mtHSP60/HSP10 promote protein folding upon import.^29, 30^ Mitochondrial chaperones can also be mobilized under stress or in response to mitochondrial protein aggregates.^29^ Indeed, exposure of islets to the saturated fatty acid palmitate to elicit lipotoxicity has been shown to upregulate mtHSP70 expression.^31^ Mitochondrial proteases not only degrade misfolded or damaged mitochondrial proteins but also have regulatory functions beyond protein clearance, including electron transport chain and mtDNA maintenance.^29^ Several well-known matrix proteases (such as LONP1 and CLPXP) govern mitochondrial protein quality control, and mutations within mitochondrial proteases have been linked to neurodegenerative diseases, cancer, and eye diseases.^32–34^ LONP1 is a multifunctional mitochondrial AAA+ matrix protease, which turns over misfolded mitochondrial proteins, remodels the ETC/OXPHOS system during tumorigenesis, and has been recently observed to possess chaperone-like activity by partnering together with mtHSP70.^35–37^ Further, LONP1 was found to be upregulated in human islets following lipotoxicity.^38^ However, the importance of LONP1 action within β-cells has not yet been explored.

Here, we unravel a crucial link between mitochondrial protein misfolding in T2D and the development of β-cell failure. Using quantitative proteomics, genetic animal models, high-resolution imaging, and biochemical assays, we elucidate those impairments in mitochondrial protein folding, which we observed in human islets of donors with T2D, elicits β-cell apoptosis, ultimately leading to loss of β cell mass and hyperglycemia. Proteomics studies reveal that insoluble/aggregated proteins in human islets of donors with T2D surprisingly more closely resembled mitochondrial rather than ER protein misfolding. Importantly, loss of LONP1, whose expression is reduced in β cells of donors with T2D, induces mitochondrial protein misfolding, thus leading to mitochondrial structural and respiratory defects, oxidative stress, DNA damage, and impairments in electron transport chain assembly. Further, we employ pharmacologic and genetically-encoded anti-oxidants to demonstrate that mitochondrial protein misfolding, and not oxidative stress, primarily contributes to β-cell death following LONP1-deficiency. Moreover, our results support that LONP1-mtHSP70 chaperone activity, and not LONP1 protease activity, prevent mitochondrial protein misfolding to promote cell survival and protect against glucolipotoxicity in mouse and human β-cells. Thus, our results illustrate the importance of mitochondrial protein folding mediated by LONP1 that may be vital to overcome β-cell loss in T2D.

## RESULTS

### Insoluble mitochondrial proteins are enriched in human islets from donors with T2D

We initially took advantage of a validated biochemical approach to evaluate protein solubility in cells or tissues, as insoluble proteins will include misfolded proteins or aggregates.^36, 39, 40^ Briefly, we collected human islets isolated from donors with or without T2D and lysed islets with a buffer containing 1% Triton X-100 ^36, 39, 40^, followed by centrifugation to isolate detergent soluble and insoluble protein fractions. We then applied unbiased quantitative proteomics approaches in both fractions by tandem mass tag (TMT)-labeling and liquid chromatography-mass spectrometry (TMT-MS; Figure 1A).

**Figure 1.**
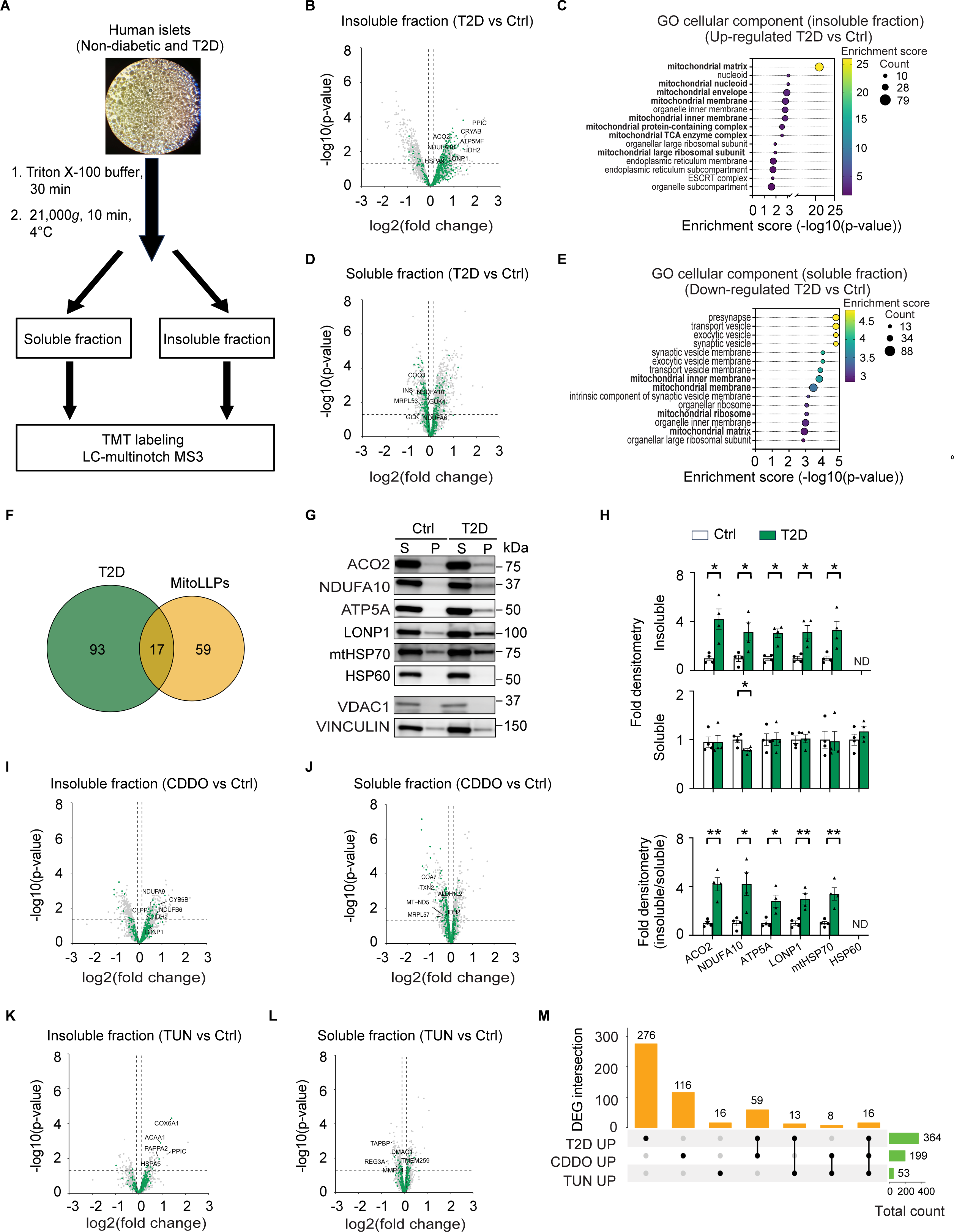
Insoluble mitochondrial proteins are enriched in human islets from donors with type 2 diabetes. (A) Schematic of Triton X-100 extraction approach to quantitatively examine solubility of proteins extracted from human islets donors that were non-diabetic or with type 2 diabetes (T2D) by tandem mass tag-labeling mass spectrometry (TMT-MS). (B) Volcano plot of differentially expressed insoluble proteins from human islet donors with T2D compared to non-diabetic controls determined by -log10 *p-*value > 1.3 and log2 fold change > 0.1. Mitochondrial proteins (curated from MitoCarta 3.0) are highlighted in green. n = 4 independent islet donors/group. (C) Gene ontology (GO) cellular component analysis of significantly upregulated insoluble proteins from human islet donors with T2D compared to non-diabetic controls. n = 4 independent islet donors/group. (D) Volcano plot of differentially expressed soluble proteins from human islet donors with T2D compared to non-diabetic controls determined by -log10 *p-*value > 1.3 and log2 fold change < -0.1. Mitochondrial proteins highlighted in green. n = 4 independent islet donors/group. (E) GO cellular component analysis of significantly downregulated soluble proteins from human islet donors with T2D compared to non-diabetic controls. n = 4 independent islet donors/group. (F) Venn diagram displaying overlap between differentially enriched mitochondrial proteins from the insoluble fraction of human islets from donors with T2D and experimentally validated mitochondrial long-lived proteins.^49^ (G) Expression of selected mitochondrial proteins by Western blot (WB) of soluble and insoluble fractions of human islets isolated from donors with T2D and non-diabetic control donors. Representative image of 4 independent human islet donors/group. VDAC1 serves as a soluble mitochondrial protein loading control. VINCULIN serves as a loading control for both soluble and insoluble fractions. (H) Quantification of protein expression by densitometry from studies in Figure 1G. Top - densitometry (fold change compared to control) of soluble and insoluble protein expression normalized to VINCULIN. Bottom - fold densitometry of insoluble/soluble protein ratio. n = 4 independent human islet donors/group. S, soluble fraction; P, insoluble fraction. Data are presented as mean ± SEM. *p < 0.05 by two-tailed Student’s *t* test. (I and J) Volcano plots for differentially expressed proteins in the insoluble (I) and soluble (J) fractions of human islets isolated from donors without diabetes exposed to 1 μM CDDO or vehicle control for 24 h. Mitochondrial proteins highlighted in green. n = 4 independent islet donors/group. (K and L) Volcano plots for differentially expressed proteins in the insoluble (K) and soluble (L) fractions of human islet donors without diabetes exposed to 1 μg/mL tunicamycin (TUN) or vehicle control for 24 h. Mitochondrial proteins highlighted in green. n = 4 independent islet donors/group. (M) UpSet blot visualizing the intersections in insoluble protein enrichment among T2D, CDDO and TUN groups.

Evaluation of our proteomics data revealed 364 proteins that were differentially enriched in the insoluble fraction of islets of T2D donors (Figure 1B). Insoluble proteins in islets of T2D donors included proteins with expected functions, such as peptidyl-prolyl cis-trans isomerase C (PPIC) ^41^, which maintains ER redox homeostasis, and α-B-crystallin (CryAB) ^42, 43^, which functions as a chaperone capable of binding misfolded proteins and toxic amyloid aggregates (Figure 1B). Of note, insulin and IAPP were increased in the insoluble fraction of islet donors with T2D yet did not reach statistical significance. Surprisingly, gene ontology (GO) analysis of differentially enriched proteins in the insoluble fraction of islets of donors with T2D revealed a high frequency of proteins localized to the mitochondria, with a striking enrichment of mitochondrial matrix proteins (Figure 1B-C). We next overlaid insoluble proteins on MitoCarta 3.0 ^44^, which contains a compendium of proteins with high confidence of localization to the mitochondria, and again confirmed the high frequency of mitochondrial proteins within the insoluble fraction of islets from donors with T2D (110 of 364 differentially enriched proteins; Figure 1B - green). Many of these enriched insoluble proteins were associated with mitochondrial gene expression, protein translation, and oxidative metabolism (Figure S1A-B). Within the soluble fraction of islets from donors with T2D, we observed a reduction of key β-cell proteins including insulin (INS) and glucokinase (GCK; Figure 1D).^45^ GO analysis of proteins differentially expressed in the soluble fraction of islets from donors with T2D included diminished proteins associated with vesicular transport and secretion, possibly related to reductions in insulin and its secretory granules, as well as increases in RNA binding, processing, and splicing, possibly related to increases in alternative splicing reported in stressed islets in diabetes (Figures 1E and S1C-D).^46–48^ Importantly, numerous mitochondrial proteins were reduced in the soluble fraction and reflected proteins that were enriched in the insoluble fraction, again localized to the mitochondrial matrix and ribosome, and related to mitochondrial translation and oxidative metabolism (Figures 1B-E, S1A-B, and S2A-B). The parallel reductions in soluble mitochondrial matrix, ribosomal, and oxidative metabolism components together with increases in these components in the insoluble fraction of islets of donors with T2D indicate an unexpected and remarkable shift in mitochondrial protein solubility in T2D, which could be attributed to mitochondrial protein misfolding.

Mitochondrial function relies on efficient proteome renewal. A recent study revealed an unusual longevity for a subset of mitochondrial proteins, including the cristae subcompartment.^49^ Mitochondrial long-lived proteins (MitoLLPs) are co-preserved within OXPHOS complexes with limited subunit exchange throughout their lifetimes, which could raise their susceptibility to aggregation. This led us to question if insoluble mitochondrial proteins in human islets of donors with T2D are long-lived or short lived, newly synthesized or imported proteins. Thus, we compared the 110 insoluble mitochondrial proteins enriched in islets of donors with T2D with a list of 76 experimentally validated MitoLLPs and observed 17 of these proteins were indeed long-lived. The observation of both insoluble short and long-lived mitochondrial proteins in islets of donors with T2D could be suggestive of a defect in mitochondrial proteostasis affecting mitochondrial resident proteins in T2D, rather than solely abnormalities in the import of newly synthesized proteins (Figure 1F).

Defects in mitochondrial proteostasis often lead to the recruitment of mitochondrial chaperones and proteases to respond to an accumulation of misfolded proteins or aggregates.^29^ Our proteomics data led us to evaluate the presence of mitochondrial proteases and chaperones in the insoluble fraction of human islets of donors with T2D. Indeed, we observed significant enrichment of the mitochondrial matrix proteases LONP1 and CLPX and chaperone HSPA9 (also known as mtHSP70) in the insoluble fraction of T2D islets (Figure 1B and S2A). Mitochondrial proteases and chaperones are also crucial for maintenance of the ETC/OXPHOS machinery. Consistent with our proteomics studies, we validated those proteins in the ETC/OXPHOS system (ACO2, NDUFA10, ATP5A) as well as LONP1 and mtHSP70 were significantly enriched in the insoluble fraction of human islets of T2D donors by western blot (WB), while no accumulation of the mitochondrial chaperone HSP60 was observed (Figure 1G-H). Notably, these ETC/OXPHOS proteins and mitochondrial chaperones/proteases were not increased in the soluble fraction of islets of T2D donors, suggesting that the accumulation of these insoluble proteins was unrelated to increased overall protein expression (Figure 1G-H).

To begin to characterize if the changes in protein solubility in T2D were similar to signatures of protein misfolding in the endoplasmic reticulum (ER) or mitochondria, we next exposed human islets from non-diabetic donors to pharmacologic agents known to elicit ER or mitochondrial protein misfolding. Following exposure of human islets to the synthetic triterpenoid CDDO, an inhibitor of LONP1 to impair mitochondrial proteostasis, or tunicamycin (TUN), a potent inhibitor of N-linked glycosylation of proteins within the ER to elicit ER protein misfolding, we again determined protein solubility by collecting soluble and insoluble fractions followed by TMT-MS.^36, 50^ As expected, both CDDO and TUN induced profound changes in protein solubility in human islets, with CDDO eliciting a robust change in mitochondrial protein solubility (Figure 1I-L). These enriched insoluble mitochondrial proteins included mitochondrial matrix proteins (CLPP and IDH2), mitochondrial membrane proteins (CYB5B), and OXPHOS subunits (NDUFB6 and NDUFA9), while reduced soluble proteins included ETC/OXPHOS system proteins and mitochondrial ribosome subunits (Figures 1I-J and S3A). As expected, TUN exposed human islets developed an increase in insoluble proteins related to ER protein misfolding or the ER unfolded protein response, such as PPIC and the ER chaperone HSPA5 (also known as BIP), with decreases in ER luminal proteins, the ERAD component membrane (TMEM259), and glycosylated proteins including TAP-associated glycoprotein (TAPBP) and matrix metallopeptidase 14 (MMP14; Figures 1K-L and S3B). Despite the well-known communication between the ER and mitochondria, we observed few changes in mitochondrial protein insolubility following TUN exposure in human islets (Figure 1K-L). On the other hand, we observed increases in ACO2, NDUFA10, ATP5A, as well as LONP1 and mtHSP70 in the insoluble fraction of CDDO-exposed human islets without accumulation of these mitochondrial proteins in the insoluble fraction of TUN-exposed islets (Figure S3C-D), suggesting a similar response of ETC/OXPHOS and mitochondrial protease/chaperone solubility in human islets in T2D and following CDDO exposure. To further visualize the intersections in insoluble protein enrichment in human islets in T2D compared to induction of mitochondrial or ER protein misfolding, we generated an UpSet plot among the T2D, CDDO, and TUN groups. Surprisingly, human islets from donors with T2D had a greater than four-fold increase in intersections of differently enriched insoluble proteins with islets following CDDO-exposure as opposed to TUN exposure (Figure 1M). Together, these data support that the robust change in mitochondrial protein solubility observed in human islets in T2D resembles, at least in part, a signature of mitochondrial protein misfolding.

### *LONP1* expression is reduced in β-cells in T2D and β-cell LONP1 deficiency results in hyperglycemia and increased β-cell apoptosis

To begin to understand the stark increase in insoluble proteins within the mitochondrial matrix in human islets of T2D donors, we profiled expression of mitochondrial matrix chaperones and proteases given their vital importance to mitochondrial proteostasis. Given previous observations of an induction of *LONP1* and *mtHSP70* mRNA expression following lipotoxicity in human and rodent islets ^31, 38^ as well as our findings of a signature of mitochondrial protein misfolding in islets of donors with T2D above, we expected to observe an induction of these genes in β-cells in T2D. Utilizing single cell RNA sequencing data from human islet donors with or without T2D (Bandesh, Motakis, Nargund, et al., in preparation), we surprisingly observed no significant increases in expression of mitochondrial matrix proteases or chaperone genes in the β-cells of islet donors with T2D (Figure 2A). Moreover, we also observed a modest, yet significant downregulation of *LONP1* within β-cells of islet donors with T2D (Figure 2A). Importantly, this effect appeared to be β-cell specific, as we did not find reduced *LONP1* expression within α-cells in T2D (Figure S4A-B). Our observations of reduced β-cell *LONP1* expression in T2D together with increases in matrix protein insolubility and aggregation in human islets of T2D donors, similar to that of LONP1 inhibition, led us to query the importance of LONP1 to β-cell health and function *in vivo*. Thus, we generated mice bearing β-cell specific deletion of *LonP1* (*LonP1*^loxP/loxP^*; Ins1*^Cre^, hereafter known as β-LonP1^KO^; Figure S2A). *LonP1*^loxP^ mice have normal glucose tolerance and body weight prior to Cre-mediated recombination as were *Ins1*^Cre^ mice per our previous studies and were thus combined as controls.^17, 51^ β-LonP1^KO^ mice exhibited an 80% reduction of LONP1 protein expression in islets compared to littermate controls (Figure 2B).

**Figure 2.**
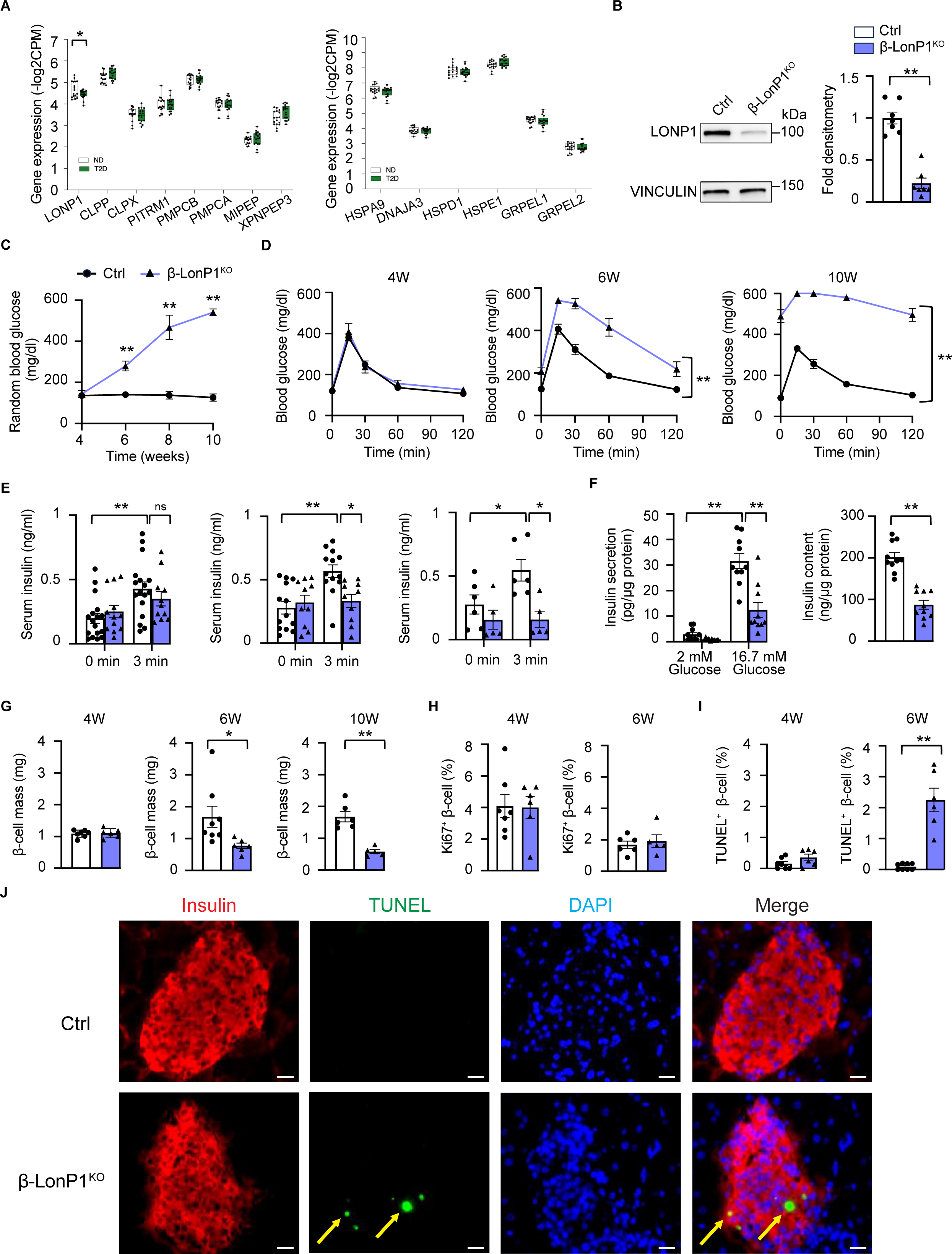
Pancreatic β-cell specific LonP1 deficiency leads to hyperglycemia due to β-cell apoptosis and loss of β-cell mass. (A) Pseudobulk gene expression data, presented as log2 counts per million reads (CPM), of mitochondrial matrix proteases and chaperones from β-cells of human islet donors with or without T2D by single cell RNA sequencing. n = 17 non-diabetic donors, n = 17 donors with T2D. *p < 0.05 by both unpaired Student’s two-tailed t-test and FDR < 5% for multiple testing correction. (B) Left - expression of LONP1 by WB in islets isolated from 4-6-week-old control (Ctrl) and β- LonP1^KO^ mice. VINCULIN serves as a loading control. Right - LONP1 protein densitometry (normalized to VINCULIN). n = 7 mice/group. **p < 0.01 by two-tailed Student’s *t* test. (C) Random blood glucose concentrations from Ctrl and β-LonP1^KO^ mice measured between age 4-10 weeks. n = 12-15/group, **p < 0.01 by one-way ANOVA followed by Tukey’s multiple comparisons test. (D) Blood glucose concentrations measured during IPGTT from Ctrl and β-LonP1^KO^ littermates at age 4-weeks, 6-weeks, and 10 weeks. n = 5-14/group. **p < 0.01 by two-way ANOVA followed by Tukey’s multiple comparisons test. (E) Serum insulin concentrations measured following *in vivo* glucose stimulation from Ctrl and β-LonP1^KO^ mice at age 4-weeks, 6-weeks, and 10-weeks. n = 5-17/group; *p < 0.05, **p < 0.01 by one-way ANOVA followed by Tukey’s multiple comparisons test. (F) Left - glucose-stimulated insulin secretion following static incubation in 2 mM and 16.7 mM glucose, performed in isolated islets of 6-week-old Ctrl and β-LonP1^KO^ littermates. Right - islet insulin content measured in isolated islets of 6-week-old Ctrl and β-LonP1^KO^ littermates. n = 10/group; **p < 0.01 by one-way ANOVA followed by Tukey’s multiple comparisons test. (G) Pancreatic β-cell mass measured in Ctrl and β-LonP1^KO^ littermates at age 4-weeks, 6-weeks, and 10-weeks. n = 5-8/group; *p < 0.05, **p < 0.01 by two-tailed Student’s *t* test. (H) Quantification of β-cell replication by Ki67 and insulin immunostaining from pancreatic sections of 4 and 6-week-old Ctrl and β-LonP1^KO^ littermates. n = 5-6/group. (I) Quantification of β-cell death by TUNEL and insulin immunostaining from pancreatic sections of 4 and 6-week-old Ctrl and β-LonP1^KO^ littermates. n = 6-8/group. **p < 0.01 by two-tailed Student’s *t* test. (J) Representative immunofluorescence image (n = 6-8/group) from pancreatic sections of 6-week-old Ctrl and β- LonP1^KO^ littermates stained for insulin (red), TUNEL (green), and DAPI (DNA - blue). Scale bar, 50 μm. Yellow arrows demarcate insulin^+^ TUNEL^+^ cells. All data within the figure are presented as mean ± SEM.

To examine the physiologic consequences of LONP1-deficiency in β-cells, we first measured random blood glucose concentrations in β-LonP1^KO^ mice and littermate controls, observing that while β-LonP1^KO^ mice were normoglycemic shortly after weaning, they developed progressive hyperglycemia beginning at 6-weeks of age (Figure 2C). Similarly, β-LonP1^KO^ mice exhibited normal glucose tolerance during an intraperitoneal glucose tolerance test (IPGTT) at 4 weeks of age, with significant and progressive glucose intolerance at 6-weeks and 10-weeks of age (Figure 2D). We also did not observe differences in body weight or peripheral insulin sensitivity between the groups (Figure S5B-C). Further, circulating insulin concentrations following an IP glucose challenge *in vivo* that progressively decreased in β-LonP1^KO^ mice with age (Figure 2E). We next assessed glucose-stimulated insulin secretion (GSIS) and insulin content in isolated islets of 6-week-old β-LonP1^KO^ mice, again observing a significant reduction compared to littermate controls (Figure 2F). We next evaluated GSIS as a fraction of total insulin content and did not identify differences between the groups (Figure S5D), suggesting reductions in β-cell insulin release could be a consequence of reduced insulin content.

The progressive reductions in glucose tolerance and insulin release with age, coupled to reduced islet insulin content, led us to question whether hyperglycemia observed in β-LonP1^KO^ mice was due to a decline in pancreatic β-cell mass. In line with our physiologic observations, β- cell mass was unchanged at 4-weeks and was significantly diminished in β-LonP1^KO^ mice by 6- and 10-weeks of age (Figure 2G). Pancreatic β-cell mass is maintained by a balance of β-cell proliferation, apoptosis, and dedifferentiation/loss of identity.^52–55^ To explore the etiology of reduced β-cell mass, we assessed markers of β-cell proliferation and apoptosis. We first assessed β-cell proliferation, observing no significant differences in β-LonP1^KO^ mice at both 4- and 6-weeks of age (Figure 2H). We next examined β-cell survival by terminal deoxynucleotidyl transferase dUTP nick end labeling (TUNEL) in pancreatic sections of control and β-LonP1^KO^ mice, noting a robust increase in the percentage of TUNEL^+^ β-cells in 6-week-old β-LonP1^KO^ mice yet no significant differences between β-LonP1^KO^ mice and littermate controls at 4-weeks of age (Figures 2I-J). As a complementary approach to assess cell survival, we evaluated apoptosis following detection of cytoplasmic histone-bound DNA fragments by ELISA in isolated islets of 6-week-old β-LonP1^KO^ mice and control littermates, which again revealed an increase in apoptosis in LONP1- deficient β-cells (Figure S5E). In addition, we did not observe reductions in the β-cell maturity marker Urocortin 3 or increases in ALDH1A3, a marker of β-cell dedifferentiation^56^, in LONP1-deficient β-cells (Figure S5F). Taken together, LONP1 deficiency leads to progressive hyperglycemia and β-cell failure related to impaired β-cell survival and loss of β-cell mass.

To ensure the physiologic and histologic changes we observed were not attributable to developmental defects, we generated inducible β-cell specific LONP1 knockout animals by intercrossing our mice bearing the LonP1 conditional allele with the tamoxifen-inducible *MIP1*-CreERT strain (*LonP1*^loxP/loxP^; *MIP1*-Cre^ERT^, hereafter known as iβ-LonP1^KO^mice). Following tamoxifen (TM)-mediated recombination at 8 weeks of age, iβ-LonP1^KO^mice exhibited glucose intolerance and reduced glucose-stimulated insulin release *in vivo* when compared to both TM-treated *MIP1*-Cre^ERT^ and vehicle-treated *LonP1*^loxP/loxP^; *MIP1*-Cre^ERT^ controls (Figures S6A-D). Further, we observed that iβ-LonP1^KO^ mice developed reductions in β-cell mass and survival without changes in β-cell replication, phenocopying constitutive β-cell LonP1 knockouts (Figure S6E-G). These data indicate that LonP1 is vital for the preservation of post-natal β-cell mass and survival that is unrelated to regulation of β-cell development.

### β-cell LONP1 deficiency results in mitochondrial protein misfolding

We next examined the specific consequences of LONP1 deficiency on β-cell mitochondrial function, structure, mass, and protein folding. We first measured mitochondrial respiration in isolated islets from 6-week-old β-LonP1^KO^ mice, observing a potent reduction in glucose-stimulated oxygen consumption compared to littermate control islets (Figure 3A). We next used transmission electron microscopy (TEM) and high-resolution three-dimensional (3D) deconvolution immunofluorescence imaging to examine mitochondrial ultrastructure and morphology as well as networking, respectively. We observed that mitochondria within β-cells from β-LonP1^KO^ mice developed severely distorted cristae, increased area, as well as alterations in matrix density which could be suggestive of an accumulation of protein aggregates by TEM (Figure 3B-C). Consistent with the observations from TEM, measures of mitochondrial morphology and networking analyzed from high-resolution 3D deconvolution imaging also revealed increases in mitochondrial volume and surface area and reductions in sphericity in LonP1-deficient β-cells, without overt defects in networking (Figure S7A-B).

**Figure 3.**
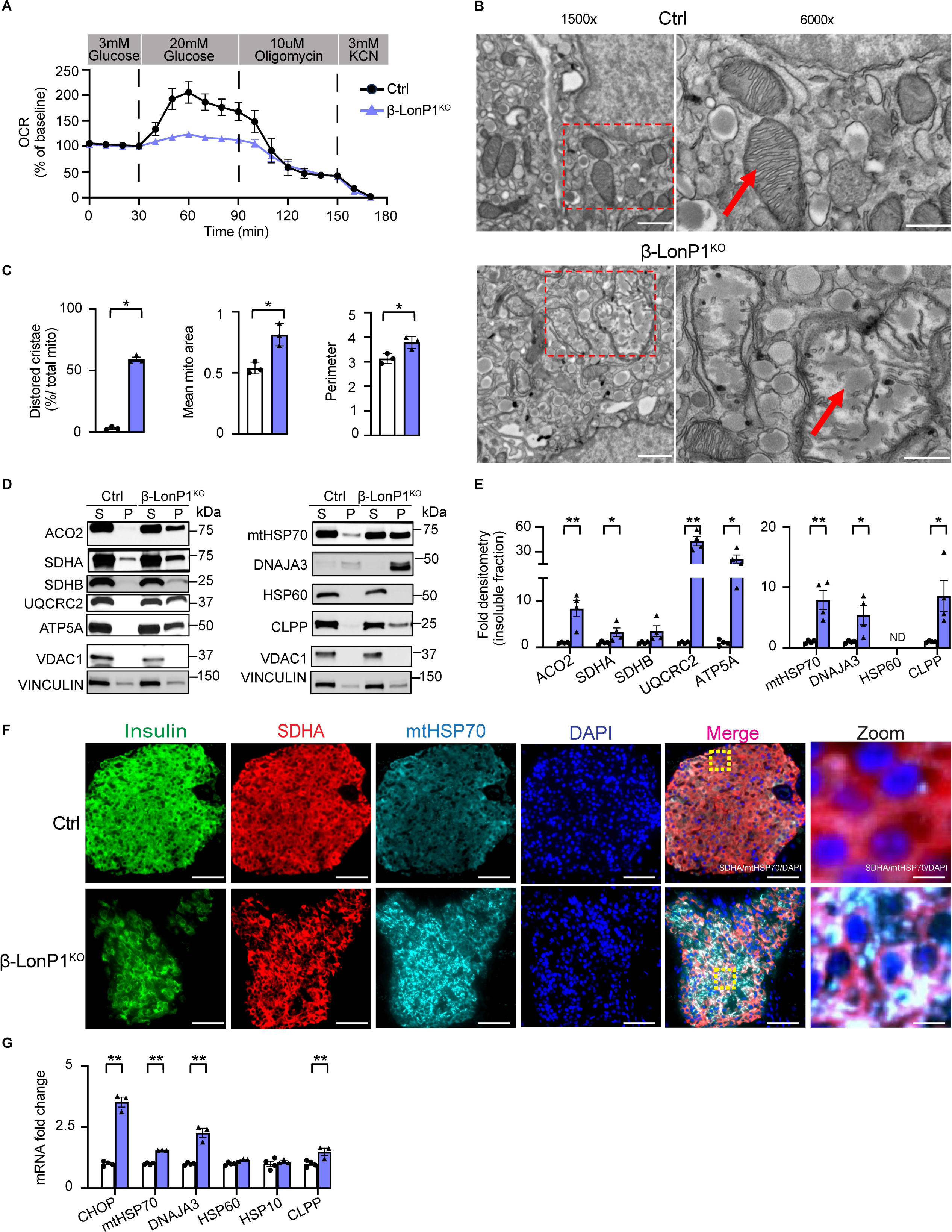
Loss of LONP1 results in impaired mitochondrial respiration, accumulation of misfolded mitochondrial proteins, and activation of the UPR^mt^ in β-cells. (A) Oxygen consumption rate (OCR) measured following exposure to 3 mM glucose, 20 mM glucose, 10 μM oligomycin and 3 mM KCN in isolated islets from 6-week-old littermate control (Ctrl) and β-LonP1^KO^ mice by a BaroFuse respirometry instrument. n = 3 mice/group. (B) Representative transmission electron microscopy (TEM) images from β-cells of 6-week-old Ctrl and β-LonP1^KO^ mice. Scale bar (left), 2 μm; Scale bar (right), 500 nm. Dotted red rectangle on left lower power images highlights focused area of mitochondria on higher power images on the right. Red arrow denotes mitochondria. n = 3 mice/group. (C) Quantification of TEM images of mitochondria (∼100 independent mitochondria scored/animal) with distorted cristae, mitochondrial area, and mitochondrial perimeter in β-cells of 6-week-old Ctrl and β-LonP1^KO^ mice. n = 3/group. *p < 0.05 by two-tailed Student’s *t* test. (D) Expression of mitochondrial metabolic proteins (left) and mitochondrial matrix chaperones and protease (right) by WB from the soluble and insoluble fractions of isolated islets from 6-week-old Ctrl and β-LonP1^KO^ mice. Representative image of 4 independent mice/group. VDAC1 serves as a soluble mitochondrial protein loading control. VINCULIN serves as a loading control for both soluble and insoluble fractions. S, soluble fraction; P, insoluble fraction. (E) Quantification of fractions of mitochondrial insoluble proteins (normalized to VINCULIN) by densitometry from studies in Figure 3D. n = 4/group. *p < 0.05 by two-tailed Student’s *t* test. (F) Representative immunofluorescence image (n = 4/group) depicting mtHSP70 expression and localization from pancreatic sections of 6-week-old Ctrl and β-LonP1^KO^ littermates, stained for insulin (green), SDHA (red), mtHSP70 (cyan) and DAPI (DNA - blue). Scale bar (Insulin, SDHA, mtHSP70, DAPI, Merge), 50 μm; Scale bar (zoom), 6.25 μm. Yellow boxes within merged image are visualized at higher magnification (Zoom - far right). White signal denotes co-localized SDHA/mtHSP70 structures within insulin^+^ cells. (G) Quantitative RT-PCR of markers of the mitochondrial unfolded protein response (UPR^mt^), normalized to *HPRT* expression, from RNA isolated from 6-week-old Ctrl and β-LonP1^KO^ islets. n = 3-4/group. *p < 0.05, **p < 0.01 by two-tailed Student’s *t* test. All data in figure are presented as mean ± SEM.

Given the known role for LONP1 as a protease with chaperone-like activity as well as reduced mitochondrial respiration and visualization of possible mitochondrial protein aggregates by TEM, we next examined whether mitochondrial protein misfolding developed in islets of β- LonP1^KO^ mice. Indeed, we observed increases in several insoluble proteins in the ETC/OXPHOS system in islets isolated from 6-week-old β-LonP1^KO^ mice (Figure 3D-E). We also observed an accumulation of mtHSP70 and its co-chaperone DNAJA3 in the insoluble fraction of 6-week-old β-LonP1^KO^ islets, as well as the mitochondrial protease CLPP (Figure 3D-E), suggestive of a response to increases in mitochondrial protein misfolding. Further, we visualized mtHSP70 expression and localization in pancreatic sections of 6-week-old control and β-LonP1^KO^ mice. We observed that mtHSP70 localized within β-cell mitochondria and LONP1-deficient β-cells developed an increase in intense and punctate mtHSP70-stained areas, possibly also indicative of the presence of mitochondrial protein aggregates (Figure 3F). Together, these results suggest that mitochondrial protein misfolding occurs in β-LonP1^KO^ islets.

The mitochondrial unfolded protein response (UPR^mt^) safeguards mitochondria from proteotoxic damage by transcriptional induction of mitochondrial chaperones and proteases to promote protein folding and eliminate misfolded proteins.^57, 58^ Therefore, we evaluated the UPR^mt^ in islets of 6-week-old β-LonP1^KO^mice and littermate controls. We observed increases in mRNA expression of several UPR^mt^ markers (Figure 3G), consistent with a transcriptional response to misfolded mitochondrial proteins in LONP1-deficient β-cells. However, a robust upregulation in mRNA expression of ER stress markers was not observed (Figure S7C).

Notably, LONP1 deficiency has been shown to regulate ETC/OXPHOS complex stability. ^32, 35^ We thus examined expression of subunits of all 5 OXPHOS complexes as well as the outer mitochondrial membrane protein TOM20 (a common marker of mitochondrial mass) by WB in islets of littermate control and β-LonP1^KO^ mice. Interestingly, a variety of changes in OXPHOS subunits were observed at 4 weeks of age in islets of β-LonP1^KO^ mice, including decreases of the complex I subunit NDUFB8 and complex IV MTCO1, while the complex III subunit UQCRC2 was increased (Figure S7D-E). At 6-weeks of age, however, β-LonP1^KO^ islets developed significant reductions of all subunits of all 5 OXPHOS complexes as well as TOM20, suggestive of a reduction in mitochondrial mass (Figure S7D-E). We next assessed whether mitochondrial mass was altered following loss of LONP1 in β-cells by several other complementary approaches. We observed a slight reduction in mtDNA content in 4-week-old β-LonP1^KO^ mice that significantly declined 6-weeks of age (Figure S7F). Further, citrate synthase activity was significantly reduced in islets of 6-week-old β-LonP1^KO^mice compared to littermate controls, while no difference was observed at 4-weeks of age (Figure S7G). Thus, these results are suggestive of early alterations of ETC/OXPHOS stability followed by a later loss of mitochondrial mass. Taken together, our results demonstrate a key role for LONP1 in the maintenance of mitochondrial proteostasis in β-cells, which preserves mitochondrial respiration, the ETC/OXPHOS machinery, and mitochondrial mass.

### Scavenging of ROS induced by LONP1 deficiency transiently improves β-cell survival and glucose homeostasis

Our results displayed a striking impairment in β-cell survival following LONP1-deficiency, yet the etiology of β-cell apoptosis was unclear. Mitochondrial dysfunction has been broadly associated with increases in the generation of free radicals that can lead to DNA damage and cell death.^59, 60^ Further, LONP1-deficiency was recently observed to induce ROS and lead to DNA damage as a key mediator of cardiomyocyte demise.^61^ ROS is a known inducer of β-cell apoptosis and reported to be elevated in human islets of donors with T2D.^62^ Historically, β-cells have been reported to have low anti-oxidant capacity, which sensitizes them to oxidative damage, although more recent work has challenged this belief.^63, 64^ Thus, we wished to determine if the generation of oxidative stress and DNA damage may be a driver of β-cell apoptosis in β-LonP1^KO^ mice. We evaluated changes in ROS between islets of 6-week-old control and β-LonP1^KO^ mice by flow cytometry, observing an increase in ROS levels in β-LonP1^KO^ islets (Figure 4A). To evaluate DNA damage, we next evaluated the phosphorylation of γH2AX by WB, a marker of DNA double strand breaks that can be increased in the setting of oxidative stress^65^, which was elevated in the islets of β-LonP1^KO^ mice (Figure 4B).

**Figure 4.**
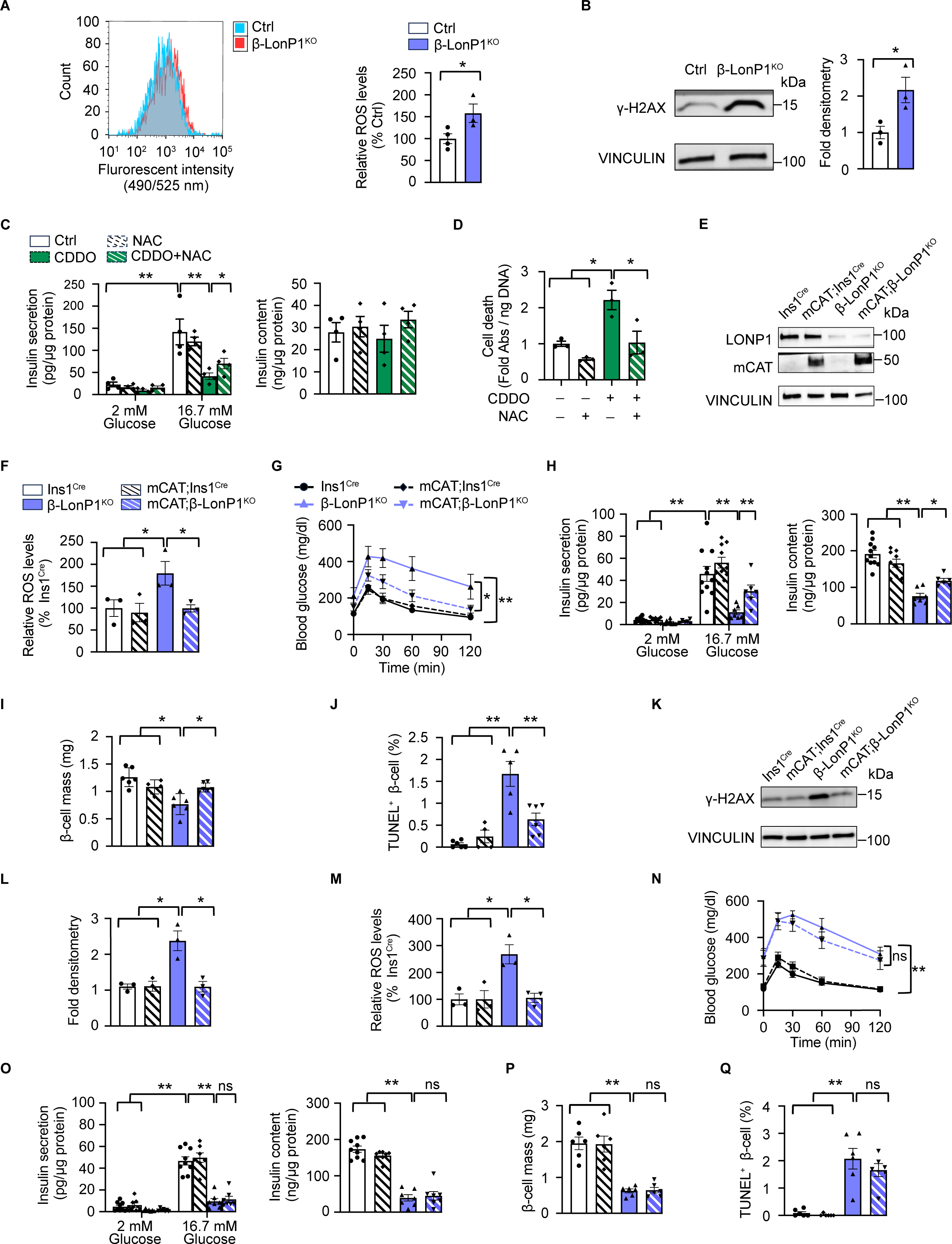
Genetic or pharmacologic free radical scavengers provide transiently improve cell survival following LONP1-deficiency in mouse and human islets. (A) Representative flow cytometry histogram demonstrating cellular reactive oxygen species (ROS) (left) and quantification of relative ROS levels in dispersed islets isolated from 6-week-old control (Ctrl) and β-LonP1^KO^ littermates. n = 3-4 mice/group; *p < 0.05 by two-tailed Student’s *t* test. (B) Left - expression of γH2AX phosphorylation by WB in islets isolated from 6-week-old control (Ctrl) and β-LonP1^KO^ mice. VINCULIN serves as a loading control. Representative of 3 mice/group. Right – phospho-γH2AX densitometry (normalized to VINCULIN). n = 3 mice/group. **p < 0.05 by two-tailed Student’s *t* test. (C) Left - glucose-stimulated insulin secretion following static incubation in 2 mM and 16.7 mM glucose, performed in human islets treated with/without 1 µM CDDO and with/without 5 mM N-acetylcysteine (NAC) for 24 h. Right - islet insulin content measured human islets treated with/without 1 µM CDDO and with/without 5 µM NAC for 24 h. n = 4 independent islet donors/group; *p < 0.05, **p < 0.01 by one-way ANOVA followed by Tukey’s multiple comparisons test. (D) Quantification of cell death by cytoplasmic histone-complexed DNA fragment ELISA (normalized to total DNA content) measured in human islets treated with/without 1 µM CDDO and with/without 5 mM NAC for 24 h. n = 4 independent islet donors/group; *p < 0.05, **p < 0.01 by one-way ANOVA followed by Tukey’s multiple comparisons test. (E) Expression of Catalase and LONP1 by WB in islets isolated from 5-7-week-old *Ins*1^Cre^, mCAT; *Ins1*^Cre^, β- LonP1^KO^, and mCAT; β-LonP1^KO^ mice. VINCULIN serves as a loading control. Representative of 3 mice/group. (F) Quantification of relative ROS levels determined by flow cytometry in isolated islets of 5-week-old *Ins*1^Cre^, mCAT; *Ins1*^Cre^, β-LonP1^KO^, and mCAT; β-LonP1^KO^ mice. n = 3 mice/group; *p < 0.05 by one-way ANOVA followed by Tukey’s multiple comparisons test. (G) Blood glucose concentrations measured during IPGTT from 5-week-old *Ins*1^Cre^, mCAT; *Ins1*^Cre^, β-LonP1^KO^, and mCAT; β-LonP1^KO^ mice. n = 6-15 mice/group; *p < 0.05, **p < 0.01 by one-way ANOVA followed by Tukey’s multiple comparisons test. (H) Left - glucose-stimulated insulin secretion following static incubation in 2 mM and 16.7 mM glucose, performed in isolated islets of 5-week-old *Ins*1^Cre^, mCAT; *Ins1*^Cre^, β-LonP1^KO^, and mCAT; β-LonP1^KO^ mice. Right - islet insulin content measured in isolated islets of 5-week-old *Ins*1^Cre^, mCAT; *Ins1*^Cre^, β-LonP1^KO^, and mCAT; β-LonP1^KO^ mice. n = 6-11 mice/group; **p<0.01 by one-way ANOVA followed by Tukey’s multiple comparisons test. (I) Pancreatic β-cell mass measured in 5-week-old *Ins*1^Cre^, mCAT; *Ins1*^Cre^, β- LonP1^KO^, and mCAT; β-LonP1^KO^ mice. n = 4-6/group; *p < 0.05 by one-way ANOVA followed by Tukey’s multiple comparisons test. (J) Quantification of β-cell death by TUNEL and insulin immunostaining from pancreatic sections of 5-week-old *Ins*1^Cre^, mCAT; *Ins1*^Cre^, β-LonP1^KO^, and mCAT; β-LonP1^KO^. n = 4-6/group; **p < 0.01 by one-way ANOVA followed by Tukey’s multiple comparisons test. (K) Expression of phospho-γH2AX by WB in islets isolated from 5-week-old *Ins*1^Cre^, mCAT; *Ins1*^Cre^, β-LonP1^KO^, and mCAT; β-LonP1^KO^ mice. VINCULIN serves as a loading control. Representative of 3 mice/group. (L) Phospho-γH2AX densitometry (normalized to VINCULIN) for studies in Figure 4K. n = 3 mice/group. *p < 0.05 by one-way ANOVA followed by Tukey’s multiple comparisons test. (M) Quantification of relative ROS production in islets of 7- week-old *Ins*1^Cre^, mCAT; *Ins1*^Cre^, β-LonP1^KO^ and mCAT; β-LonP1^KO^ mice. n = 3 mice/group; *p < 0.05 by one-way ANOVA followed by Tukey’s multiple comparisons test. (N) Blood glucose levels measured during IPGTT from 7-week-old *Ins*1^Cre^, mCAT; *Ins1*^Cre^, β-LonP1^KO^, mCAT and β- LonP1^KO^ mice. n = 4-11 mice/group; **p < 0.01 by one-way ANOVA followed by Tukey’s multiple comparisons test. (O) Left - glucose-stimulated insulin secretion following static incubation in 2 mM and 16.7 mM glucose, performed in isolated islets of 7-week-old *Ins*1^Cre^, mCAT; *Ins1*^Cre^, β- LonP1^KO^, and mCAT; β-LonP1^KO^ mice. Right - islet insulin content measured in isolated islets of 7-week-old *Ins*1^Cre^, mCAT; *Ins1*^Cre^, β-LonP1^KO^, and mCAT; β-LonP1^KO^ mice. n = 7-9 mice/group; **p<0.01 by one-way ANOVA followed by Tukey’s multiple comparisons test. (P) β-cell mass determined in pancreatic sections of 7-week-old *Ins*1^Cre^, mCAT; *Ins1*^Cre^, β-LonP1^KO^, and mCAT; β-LonP1^KO^ mice. n = 6 mice/ group; **p < 0.01 by one-way ANOVA followed by Tukey’s multiple comparisons test. (Q) Quantification of TUNEL staining performed in pancreatic sections of 7- week-old *Ins*1^Cre^, mCAT; *Ins1*^Cre^, β-LonP1^KO^, and mCAT; β-LonP1^KO^ mice for β-cell apoptosis. n = 6 mice/group; **p < 0.01 by one-way ANOVA followed by Tukey’s multiple comparisons test. All data in figure are presented as mean ± SEM.

To determine if ROS contributes to β-cell failure following LONP1-deficiency, we employed both pharmacologic and genetic approaches to ameliorate oxidative stress. To initially address this question in human islets, we exposed islets to the LONP1 inhibitor CDDO, observing reductions in GSIS and cell survival similar to our observations in β-LonP1^KO^ islets (Figure 4C-D). Interestingly, use of the antioxidant N-acetylcysteine (NAC) ameliorated defects in GSIS and cell survival in human islets following acute LONP1 inhibition (Figure 4C-D). To test if excess mtROS led to β-cell dysfunction following LonP1 deficiency *in vivo*, we intercrossed β-LonP1^KO^ mice (or *Ins1^Cre^* controls) with Cre-inducible mitochondrial-targeted catalase (mCAT) overexpression mice to selectively scavenge β-cell mtROS as mCAT overexpression has been shown to reduce H_2_O_2_ as well as superoxide in β-cells.^66, 67^ As expected, we found efficient expression of catalase in islets of mCAT; β-LonP1^KO^ mice (Figure 4E) with a restoration of ROS to levels similar of *Ins1^Cre^* and mCAT; *Ins1^Cre^* controls (Figure 4F). Overexpression of mCAT improved glucose intolerance, glucose-stimulated insulin release, β-cell mass, and β-cell survival following LONP1-deficiency at age 5 weeks (Figure 4G-J). In addition, overexpression of mCAT lowered phosphorylation of γH2AX following LONP1-deficiency (Figure 4K-L). The beneficial effects of mCAT overexpression in LONP1-deficient animals were short-lived, however, as mCAT overexpression did not sustain improvements in glucose tolerance, serum insulin levels, β-cell mass, and β-cell survival by 7- weeks of age, despite continued improvements in ROS levels (Figure 4M-Q). Together, these observations suggest that while reductions in ROS elicit an acute and transient protective effect on β-cell survival, oxidative stress is unlikely to be the primary mediator of β-cell demise following LONP1 deficiency.

### Mitochondrial protein misfolding precedes oxidative stress following LONP1 deficiency

Accumulating evidence indicates that an interaction between misfolded proteins and oxidative stress contributes to β-cell failure in the development of diabetes.^68, 69^ Our observations of mitochondrial protein misfolding and oxidative stress following LONP1-deficiency next led us to determine if ROS induced mitochondrial protein misfolding. We first evaluated soluble and insoluble protein fractions in human islets exposed to CDDO, observing an accumulation of ETC/OXPHOS proteins as well as the mtHsp70 chaperone machinery and LONP1 itself in the insoluble fraction (Figure 5A-B). Following NAC treatment, we did not observe improvements in mitochondrial protein misfolding in human islets following CDDO exposure (Figure 4A-B). We next evaluated mitochondrial protein misfolding in islets from β-LonP1^KO^ mice as well as LONP1-deficient animals bearing mCAT overexpression. Similar to observations in human islets, we found that β-cell mCAT overexpression in mice was unable to relieve misfolding of ETC/OXPHOS proteins or presence of the mtHSP70 machinery and CLPP in the insoluble fraction in LONP1-deficient β-cells both at 5 and 7 weeks of age (Figure 5C-D). These results suggest that β-cell mitochondrial protein misfolding following LONP1 deficiency is not a consequence of oxidative stress. These results also led us to speculate that the continued presence of mitochondrial protein misfolding in mCAT; β-LonP1^KO^ mice may override the transient protective effects of antioxidant exposure, ultimately leading to hyperglycemia and loss of β-cell mass.

**Figure 5.**
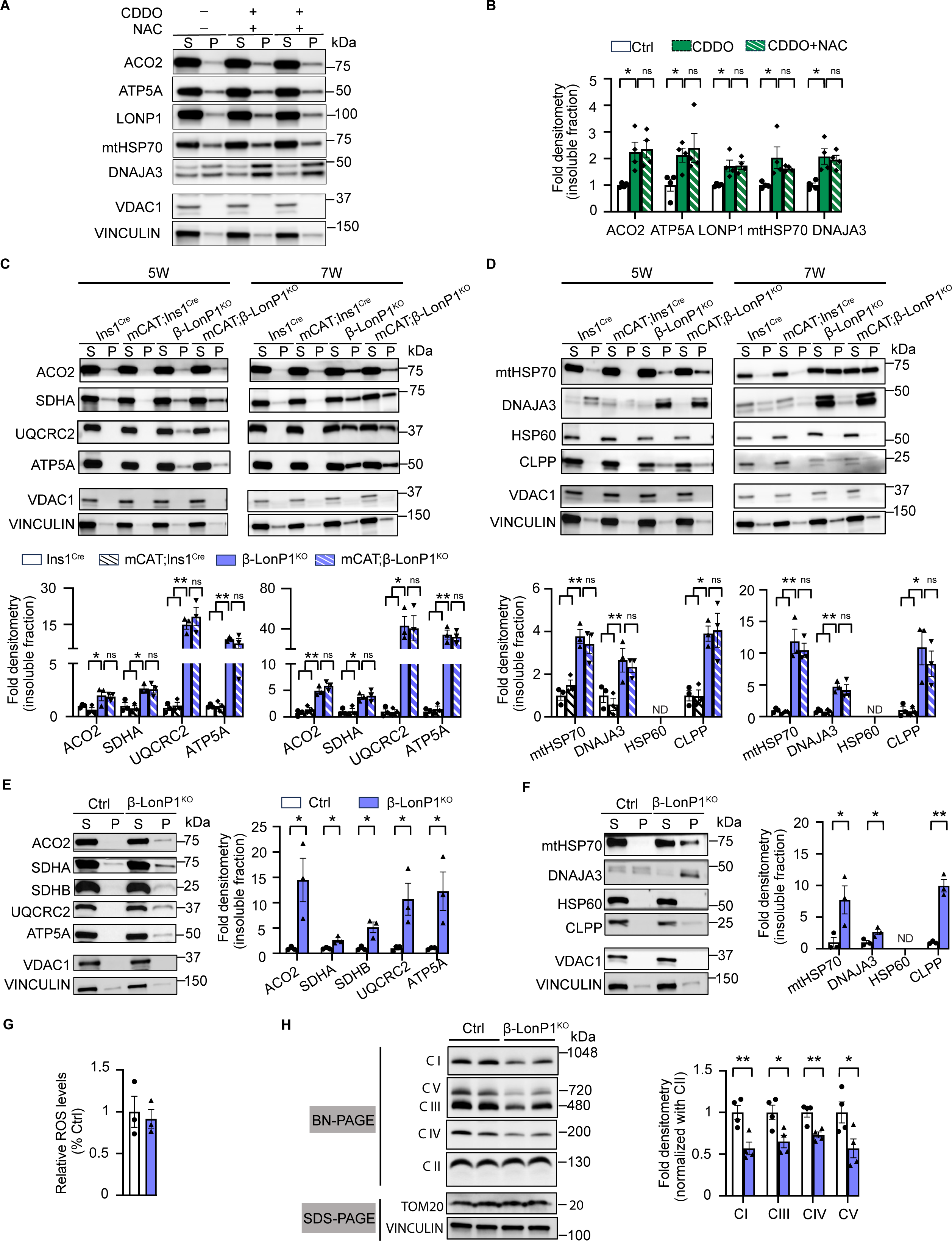
Mitochondrial protein misfolding is not ameliorated by antioxidants and precedes the appearance of oxidative stress following LONP1-deficiency. (A) Expression of selected mitochondrial metabolic proteins, matrix chaperones, and proteases by WB from the soluble and insoluble fractions of human islets exposed to DMSO (Ctrl), 1 μM CDDO, or 1 μM CDDO + 5 mM NAC for 24 h. Representative image of 4 independent human islet donors/group. VDAC1 serves as a soluble mitochondrial protein loading control. VINCULIN serves as a loading control for both soluble and insoluble fractions. S, soluble fraction; P, insoluble fraction. (B) Quantification of fractions of mitochondrial insoluble proteins (normalized to VINCULIN) by densitometry from studies in Figure 5A. n = 4 independent human islet donors/group. *p < 0.05 by one-way ANOVA followed by Tukey’s multiple comparisons test. (C) Expression of ETC/OXPHOS system proteins by WB from the soluble and insoluble fractions of isolated islets from *Ins*1^Cre^, mCAT; *Ins1*^Cre^, β-LonP1^KO^, and mCAT; β-LonP1^KO^ mice at both 5-weeks and 7-weeks of age. Representative image (upper) of 3 independent mice/group. Quantification of fractions of mitochondrial insoluble proteins (normalized to VINCULIN) by densitometry from studies is shown in graphs below representative WBs. VDAC1 serves as a soluble mitochondrial protein loading control. VINCULIN serves as a loading control for both soluble and insoluble fractions. S, soluble fraction; P, insoluble fraction. n = 3 independent mice/group. *p < 0.05, **p < 0.01 by one-way ANOVA followed by Tukey’s multiple comparisons test. (D) Expression of mitochondrial matrix chaperones and proteases by WB from the soluble and insoluble fractions of isolated islets from *Ins*1^Cre^, mCAT; *Ins1*^Cre^, β-LonP1^KO^, and mCAT; β-LonP1^KO^ mice at both 5-weeks and 7-weeks of age. Representative image (upper) of 3 independent mice/group. Quantification of fractions of mitochondrial insoluble proteins (normalized to VINCULIN) by densitometry from studies is shown in graphs below representative WBs. VDAC1 serves as a soluble mitochondrial protein loading control. VINCULIN serves as a loading control for both soluble and insoluble fractions. S, soluble fraction; P, insoluble fraction. n = 3 independent mice/group. *p < 0.05, **p < 0.01 by one-way ANOVA followed by Tukey’s multiple comparisons test. (E) Expression of ETC/OXPHOS system proteins by WB from the soluble and insoluble fractions of isolated islets of 4-week-old Ctrl and β-LonP1^KO^ littermates. Representative images (left) of 3 independent mice/group. Quantification of mitochondrial insoluble proteins (normalized to VINCULIN) by densitometry is shown in graph on right. VDAC1 serves as a soluble mitochondrial protein loading control. VINCULIN serves as a loading control for both soluble and insoluble fractions. S, soluble fraction; P, insoluble fraction. n = 3 independent mice/group. *p < 0.05 by two-tailed Student’s *t* test. (F) Expression of mitochondrial chaperones and proteases by WB from the soluble and insoluble fractions of isolated islets of 4-week-old Ctrl and β-LonP1^KO^ littermates. Representative images (left) of 3 independent mice/group. Quantification of mitochondrial insoluble proteins (normalized to VINCULIN) by densitometry is shown in graph on right. VDAC1 serves as a soluble mitochondrial protein loading control. VINCULIN serves as a loading control for both soluble and insoluble fractions. S, soluble fraction; P, insoluble fraction. n = 3 independent mice/group. *p < 0.05, **p < 0.01 by two-tailed Student’s *t* test. (G) Quantification of relative ROS production in islets of 4-week-old Ctrl and β-LonP1^KO^ littermates. n = 3 mice/group. (H) Blue native PAGE (BN-PAGE) followed by immunoblotting for OXPHOS complexes performed in isolated islets from 4-week-old Ctrl and β-LonP1^ko^ littermates. Representative image (left) of studies from n = 4/group. Quantification of complexes I, III, IV, and V (normalized to complex II) by densitometry from BN-PAGE studies is shown in graph on right. SDS-PAGE for TOM20 and VINCULIN represent additional loading controls from these samples. n = 4/group. *p < 0.05, **p < 0.01 by two-tailed Student’s *t* test. All data in figure are presented as mean ± SEM.

To better clarify the chronologic etiology of mitochondrial dysfunction leading sequentially to β-cell failure after loss of LONP1, we assessed key mitochondrial defects visible in β-LonP1^KO^ mice at 4-weeks of age. As we previously showed, no differences in glycemic control, glucose-stimulated insulin release, β-cell mass, β-cell survival, or mitochondrial mass were visible in β-LonP1^KO^ mice compared to littermate controls at this age (Figures 2C-E, 2G, 2I, and S7F-G), thus allowing us to identify mechanistic defects that are forerunners of β-cell dysfunction and not related to hyperglycemia or glucotoxicity. Interestingly, we observed evidence of mitochondrial protein misfolding in β-LonP1^KO^ islets, including increases in insoluble ETC/OXPHOS proteins as well as the mtHSP70 chaperone machinery and CLPP (Figure 5E-F). Notably, these increases in insoluble mitochondrial proteins developed following LONP1-deficiency despite no differences in islet ROS levels or oxidative DNA damage at this age (Figure 5G and S7H).

Our observations of insoluble ETC/OXPHOS subunits preceding hyperglycemia in β-LonP1^KO^ mice next led us to query if formation of fully assembled ETC complexes were impaired. Indeed, loss of key subunits of complexes I, III, and IV within β-cells have all been shown to lead to hyperglycemia.^70^ Thus, we examined ETC complexes by blue-native PAGE (BN-PAGE).^71^ Importantly, complexes I, III, IV and V were all decreased in islets of 4-week old β-LonP1^KO^ mice (Figure 5J), prior to changes in mitochondrial mass (Figure S7D-G). Taken together, these observations suggest that mitochondrial protein misfolding, and not oxidative stress, is the initial insult detected following LONP1-deficiency, which heralds the development of ETC/OXPHOS system defects, β-cell failure, and hyperglycemia.

### LONP1-mtHSP70 chaperone activity promotes β-cell survival by relieving mitochondrial protein misfolding in mouse and human islets

Due to recent studies outlining several roles for LONP1 within mitochondria^36, 37^, we next wished to clarify the specific mechanism by which LONP1 acts to prevent mitochondrial protein misfolding and maintain β-cell survival. To determine if LONP1 requires protease activity to promote β-cell survival, we expressed a protease-deficient LONP1 S855A mutant ^32, 36, 72^ or empty vector control (EV) in β-cells of islets of 5-week-old β-LonP1^KO^ mice (or littermate controls) using the recently described pseudoislet approach.^73^ Briefly, primary islets were dispersed, transduced with adenoviral particles encoding LONP1 S855A or EV under control of the rat insulin 2 promoter (Ad.*RIP2*.LONP1^S855A^ or Ad.*RIP2*.EV, respectively) to facilitate β-cell specific expression, and then re-aggregated into pseudoislets (Figure 6A). We then assessed β-cell survival 7 days after pseudoislet generation following dissociation and cytocentrifugation onto slides for TUNEL analysis. Interestingly, re-expression of the protease dead LONP1 S855A mutant significantly rescued β-cell apoptosis in β-LonP1^KO^ pseudoislets (Figure 6B-C). Given the partnership between LONP1 and mtHSP70 necessary for LONP1 chaperone-like activity^36^, we next exposed β-LonP1^KO^ pseudoislets to MKT077, an inhibitor of mtHSP70^74, 75^, to determine if the beneficial effects of LONP1 S855A re-expression were related to LONP1-mtHSP70 function. Indeed, MKT077 exposure abrogated the rescue of Ad.*RIP2*.LONP1^S855A^ on cell survival in β-LonP1^KO^ pseudoislets, suggesting that LONP1-mtHSP70 chaperone activity is necessary to maintain β-cell survival (Figure 6B-C).

**Figure 6.**
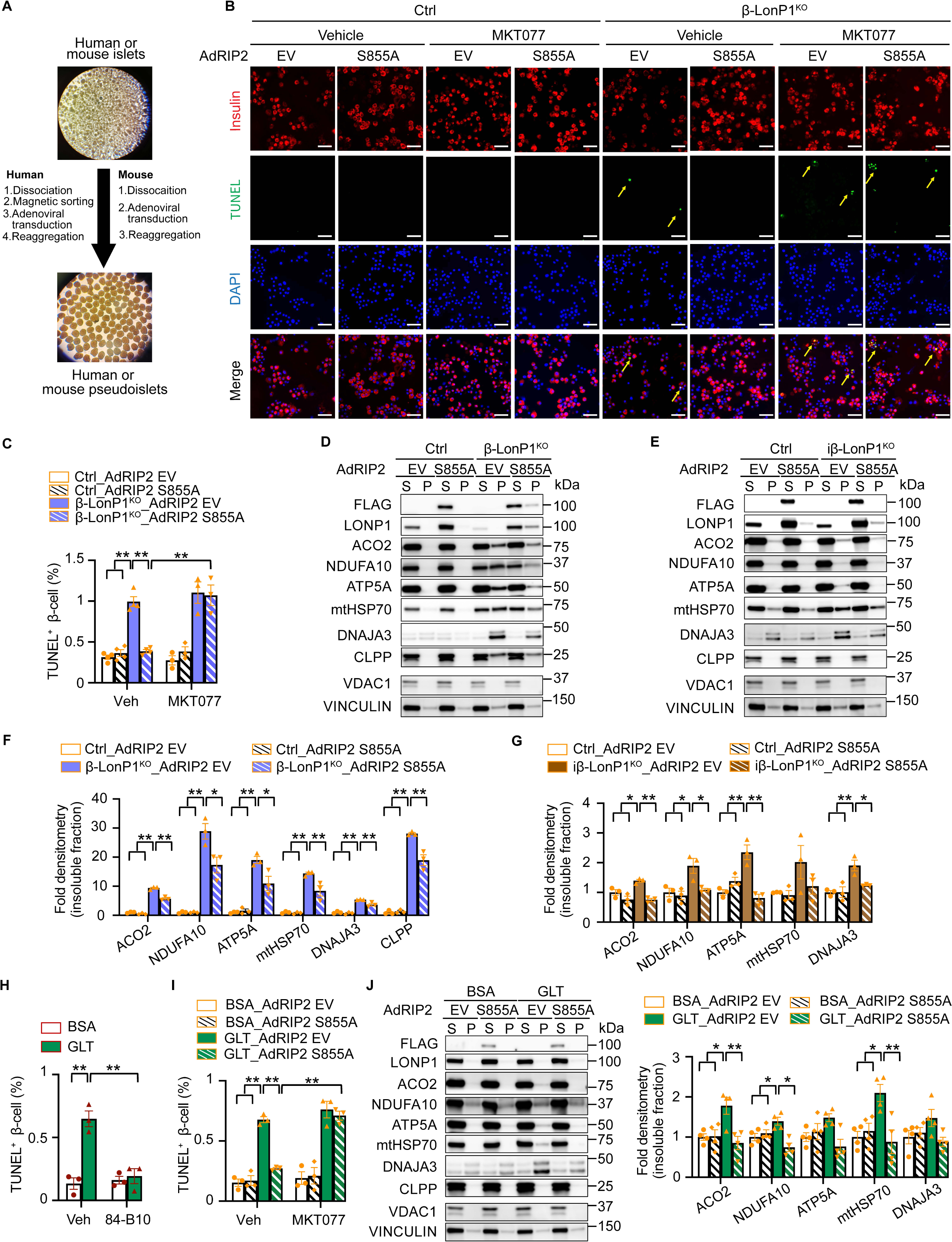
LONP1-mtHSP70 chaperone activity promotes β-cell survival by relieving mitochondrial protein misfolding. (A) Schematic for generation of mouse or human pseudoislets. (B) TUNEL staining for β-cell apoptosis performed in mouse pseudoislets dissociated for cytocentrifugation from 5-week-old control (Ctrl) and β-LonP1^KO^ littermates performed 7 days after adenoviral transduction with *RIP2*-driven empty vector (Ad *RIP2* EV) and protease-deficient LONP1 S855A mutant (Ad *RIP2* S855A), with exposure to vehicle (DMSO) or 1 μM mtHSP70 inhibitor MKT077 for the final 24 hrs. Representative images of 3-4 mice/group. Insulin (red), TUNEL (green), and DAPI (DNA - blue). Scale bar, 50 μm. Yellow arrows demarcate insulin^+^ TUNEL^+^ cells. (C) Quantification of TUNEL staining from studies in Figure 6B. n = 3-4 mice/group. **p < 0.01 by one-way ANOVA following Tukey’s multiple comparisons test. (D) Expression of mitochondrial proteins from the soluble and insoluble fractions performed in mouse pseudoislets from 5-week-old Ctrl and β-LonP1^KO^ mice 7 days after adenoviral transduction with Ad *RIP2* EV or Ad *RIP2* S855A by WB. Representative images of 3 mice/group. VDAC1 serves as a soluble mitochondrial protein loading control. VINCULIN serves as a loading control for both soluble and insoluble fractions. S, soluble fraction; P, insoluble fraction. (E) Expression of mitochondrial proteins from soluble and insoluble fractions of mouse pseudoislets generated from 8-week-old *LonP1*^loxP/loxP^; MIP-Cre^ERT^ mice 7 days after adenoviral transduction with Ad *RIP2* EV or Ad *RIP2* S855A. Pseudoislets were co-cultured with vehicle (EtOH) or 2 μM 4-hydroxytamoxifen (4-OHT), to induce recombination *in vitro* and generate experimental groups (Ctrl or iβ-LonP1^KO,^ respectively), prior to generation of soluble or insoluble fractions for WB. Representative images of 3 mice/group. VDAC1 serves as a soluble mitochondrial protein loading control. VINCULIN serves as a loading control for both soluble and insoluble fractions. S, soluble fraction; P, insoluble fraction. (F) Quantification of mitochondrial insoluble proteins (normalized to VINCULIN) from studies in Figure 6D. n = 3-4 mice/group. *p < 0.05, **p < 0.01 by one-way ANOVA followed by Tukey’s multiple comparisons test. (G) Quantification of mitochondrial insoluble proteins (normalized to VINCULIN) from studies in Figure 6E. n = 3 mice/group. *p < 0.05, **p < 0.01 by one-way ANOVA followed by Tukey’s multiple comparisons test. (H) Quantification of TUNEL staining for β-cell apoptosis in human β-cell enriched pseudoislets dissociated for cytocentrifugation following exposure to BSA or GLT (25 mM glucose + 0.5 μM palmitate) together with vehicle (DMSO) or 40 μM LONP1 activator 84-B10 for 48 hrs. n = 3 independent human islet donors/group. **p < 0.01 by one-way ANOVA followed by Tukey’s multiple comparisons test. (I) Quantification of TUNEL staining for β-cell apoptosis in human β-cell enriched pseudoislets dissociated for cytocentrifugation 7 days after Ad *RIP2* EV or Ad *RIP2* S855A transduction, followed by exposure to BSA or GLT for the final 48 hrs, as well as vehicle (DMSO) or 1 μM MKT077 for the final 24 hrs. n = 3 independent human islet donors/group. **p < 0.01 by one-way ANOVA followed by Tukey’s multiple comparisons test. (J) Expression of mitochondrial proteins from the soluble and insoluble fractions of human β-cell enriched pseudoislets 8 days after Ad *RIP2* EV or Ad *RIP2* S855A transduction, exposed to BSA or GLT for the final 72 hrs. Representative images (left) of 4 independent islet donors/group. Quantification of mitochondrial insoluble proteins by densitometry (normalized to VINCULIN) appears in graph on right. VDAC1 serves as a soluble mitochondrial protein loading control. VINCULIN serves as a loading control for both soluble and insoluble fractions. S, soluble fraction; P, insoluble fraction. n = 4 independent islet donors/group. *p < 0.05, **p < 0.01 by one-way ANOVA followed by Tukey’s multiple comparisons test. All data in figure are presented as mean ± SEM.

We next wished to determine the role of LONP1 protease activity on mitochondrial protein misfolding, observing that Ad.*RIP2*.LONP1^S855A^ transduction significantly reduced insoluble ETC/OXPHOS proteins as well as the mtHSP70 chaperone machinery and CLPP in β-LonP1^KO^ pseudoislets (Figure 6D-E). However, we only observed a partial rescue of mitochondrial protein misfolding 7 days after Ad.*RIP2*.LONP1^S855A^ transduction in islets of 5-week-old constitutive β-LonP1^KO^ mice. This led us to speculate that the prolonged burden of misfolded mitochondrial proteins could not be completely cleared within 1 week of LONP1 S855A re-expression. Thus, to elicit acute formation of misfolded mitochondrial proteins due to loss of LONP1, we next generated pseudoislets from iβ-LonP1^KO^ mice following *in vitro* recombination achieved in pseudoislets cultured in the presence of 4-hydroxytamoxifen (or vehicle control). We again observed increases in ETC/OXPHOS proteins and the mtHSP70 chaperone machinery in the insoluble fraction of iβ-LonP1^KO^ pseudoislets (Figure 6F-G). Importantly, transduction of iβ-LonP1^KO^ pseudoislets with Ad.*RIP2*.LONP1^S855A^ reversed the accumulation of misfolded mitochondrial proteins (Figure 6F-G), consistent with a key protease-independent role for LONP1 to maintain proper mitochondrial protein folding in β-cells.

Finally, to explore the relevance of LONP1 and its mechanism of action in a model of T2D in human β-cells, we assessed β-cell survival in human islets subjected to GLT. Notably, GLT has been reported to elicit β-cell toxicity, the UPR, and mitochondrial structural/functional defects ^76–, 79^, yet a role for GLT to induce mitochondrial protein misfolding, similar to our observations in human islets in T2D donors (Figure 1), has not previously been assessed. Interestingly, the LONP1 activator 84-B10 ^80^ rescued GLT-induced β-cell apoptosis in human islets (Figure 6H). To determine if LONP1 protease or LONP1-mtHSP70 chaperone activity were necessary to promote human β-cell survival and mitochondrial protein folding, we next generated human β-cell enriched pseudoislets following magnetic sorting for the β-cell marker NTPDase3 for better visualization of β-cell proteins in subsequent biochemical assays. Indeed, we confirmed that magnetically sorting for NTPDase3 yielded ∼90-95% insulin^+^ β-cells (Figure S8).^81^ Human β-cells were then transduced with Ad.*RIP2*.LONP1^S855A^ (or Ad.*RIP2*.EV control) prior to pseudoislet generation (Figure 6A). Excitingly, we observed that overexpression of LONP1 S855A rescued GLT-induced β-cell apoptosis (Figure 6I). Similar to observations in mouse islets above, the effects of LONP1 to rescue β-cell survival following GLT were dependent on LONP1-mtHSP70 activity, as the benefits of LONP1 S855A overexpression were abrogated in the presence of MKT077 exposure (Figure 6I). Interestingly, we observed that GLT exposure also elicited a signature of mitochondrial protein misfolding in human β-cells, which was reversed following overexpression of the LONP1 S855A mutant (Figure 6J). Taken together, we observe that LONP1 acts together with the chaperone mtHSP70 to improve human β-cell survival following glucolipotoxicity.

## DISCUSSION

Here, we identify impaired mitochondrial proteostasis in islets in T2D and elucidate the impact of mitochondrial protein misfolding on β-cell survival and glycemic control. We find that β-cells of T2D donors exhibit reduced expression of the matrix protease *LONP1*, and loss of *LONP1* elicits an accumulation of misfolded mitochondrial proteins that subsequently yields bioenergetic defects and oxidative stress. Intriguingly, LONP1 acts in a protease-independent manner together with the chaperone mtHSP70 to promote mitochondrial protein folding and rescue human β-cell survival following GLT to model the metabolic stress of T2D. Together, our studies position mitochondrial proteostasis as a crucial regulator of β-cell survival and provide possible new insights into mechanisms of β-cell failure in T2D.

To understand the extent of impaired proteostasis in T2D, we applied a biochemical approach to quantitative islet proteomics by identification of changes in protein solubility. We were surprised to observe a robust enrichment in insoluble proteins within the mitochondrial matrix in islets of donors with T2D. Further, we unexpectedly found that islets of donors with T2D had a signature favoring protein misfolding within the mitochondria to an increased extent than ER, positioning mitochondrial proteostasis as a new possible pathway in the pathology of islets in T2D. Recent rigorous quantitative proteomics studies on islets from donors with T2D also support the observation of defective mitochondrial energy metabolism in T2D.^21, 82^ Together with these studies, our work supports tailoring the use of future discovery studies in T2D to discern organelle or compartment-specific events that transpire to yield β-cell dysfunction. Future studies may also be expanded to include spatial single cell technologies in T2D, such as Co-Detection by Indexing (CODEX) and Imaging Mass Cytometry (IMC), to leverage high-throughput visualization of islet cell organization and composition towards the evaluation of organelle-specific events.^82–85^

Despite its primary role as a mitochondrial protease, models of LONP1 deficiency display numerous mitochondrial defects, including the accumulation of misfolded mitochondrial protein aggregates, increases in mtROS, alterations in the ETC/OXPHOS system, and reductions of mitochondrial mass ^35–37, 61^. We observed many of these effects in β-cells bearing *LONP1* deficiency and endeavored to identify an initiating cause of β-cell dysfunction and apoptosis in our models. Indeed, we observed that oxidative damage was a slight contributor to loss of β-cell mass and hyperglycemia in β-LonP1^KO^ mice; however, overexpression of a mitochondrial targeted catalase was insufficient to remedy mitochondrial protein misfolding and only briefly delayed β-cell failure. We also observed the induction of mitochondrial protein misfolding, including key ETC/OXPHOS proteins, that was associated with reduced levels of fully assembled ETC complexes that preceded ROS and reductions in mitochondrial mass. Interestingly, LONP1 deficiency has also been found to elicit autophagy/mitophagy as a compensatory measure to clear mitochondria bearing misfolded proteins, which will be of interest for future investigation given the importance of autophagy/mitophagy in β-cells.^32, 33, 86^ We observe that LONP1 acts in a protease-independent manner together with the chaperone mtHSP70 to regulate β-cell survival and prevent mitochondrial protein misfolding. These results suggest that in β-cells, prevention of mitochondrial protein misfolding by LONP1 may be vital to avert a cascade of additional mitochondrial defects.

While *LONP1* is not a known risk gene for T2D, our studies suggest connections between LONP1, mitochondrial protein misfolding, and β-cell viability of value to the understanding of T2D. A question raised from our work is not only the origin of reduced β-cell *LONP1* expression in T2D, but also the additional failure to upregulate LONP1 or mtHSP70 expression in response to mitochondrial protein misfolding, a phenomenon which had been shown to occur following exposure of human or mouse islets to lipotoxicity to model events of T2D.^31, 38^ Indeed, the lack of increased β-cell expression of matrix proteases and chaperones in T2D may additionally suggest a failed transcriptional response to misfolded mitochondrial proteins that will be enticing for future exploration. Of course, these defects may also be related to polymorphisms in key nuclear-encoded mitochondrial genes associated with T2D as *LONP1* has not been identified as a risk locus for T2D.^22–28^ We also observe evidence for mitochondrial protein misfolding and subsequent apoptosis of human β-cells following GLT that are ameliorated by LONP1 overexpression, insinuating that even relatively acute diabetogenic stimuli might begin to overwhelm the endogenous mitochondrial proteostasis machinery in β-cells of non-diabetic donors. While our early results could inspire possible future therapeutic targets related to LONP1, these studies may also insinuate limitations in the intrinsic capacity of β-cells to balance mitochondrial proteostasis in the face of metabolic stress that will be of interest for future study.

T2D is a complex polygenic and multi-organ disease driven by β-cell dysfunction and complicated by extrinsic metabolic and chronologic stress whose etiopathological mechanisms remain vexing.^87, 88^ Our studies here identify a convergence of two key β-cell defects of impaired proteostasis and mitochondrial dysfunction in T2D due to mitochondrial protein misfolding. The promising effects to improve β-cell survival by pharmacological chaperones to offset ER protein misfolding as well as proteases targeting IAPP suggest that future approaches to improve mitochondrial proteostatic responses might be beneficial during metabolic stress.^89–91^ Interestingly, activation of mitochondrial proteases and chaperones without the induction of mitochondrial damage have had promise in the extension of lifespan in *Caenorhabditis elegans*.^92^ It is thus intriguing to speculate if activation of mitochondrial proteases and chaperones to improve β-cell mitochondrial proteostasis could also improve metabolic efficiency and viability in the treatment or prevention of T2D.

## MATERIALS AND METHODS

### Genetically modified mouse lines

All mice were maintained in accordance with the University of Michigan’s Institutional Animal Care and Use Committee under specific pathogen-free conditions. Up to 5 mice were housed per cage and were maintained with *ad libitum* access to food on a 12 h light-dark cycle. All experiments were performed with both male and female mice, and results from both sexes were combined in all experimental groups. *LonP1*^loxP/loxP^ tm1a mice possessing loxP sites flanking exon 2 of the *LonP1* gene were obtained from the European Mutant Mouse Archive (EMMA). These animals were initially crossed to the *ACTB*-FLPe line to achieve deletion of the FRT-flanked Neomycin cassette ^93^, prior to generation of tissue specific knockout models. To generate β-cell-specific deletion, floxed models were crossed with *Ins1*-Cre mice from Jackson laboratories (JAX Stock No. 026801). *Ins1*^Cre^–alone and floxed-only controls (*LonP1*^loxP/loxP^ and *LonP1*^loxP/+^ mice) were phenotypically indistinguishable from each other and combined as controls (Ctrl). *Ins1*^Cre^–alone were also phenotypically indistinguishable from wild-type C57BL/6N controls, consistent with previous reports from our group and others^17, 94–96^. For inducible β-cell deletion, *MIP1*-CreERT mice (JAX 024709) were used or crossed to *LonP1*^loxP/loxP^ mice to generate iβ-LonP1^KO^ mice. 8-week-old *MIP1*-Cre^ERT^; *LonP1* ^loxP/loxP^ mice and *MIP1*-Cre^ERT^ mice were administered 100 mg/kg/d tamoxifen (TM, T5648, Sigma-Aldrich) dissolved in filtered sunflower seed oil (Sigma-Aldrich) by intraperitoneal injection for 5 consecutive days. Filtered sunflower seed oil was administered into *MIP1*-Cre^ERT^; *LonP1* ^loxP/loxP^ mice as an additional control group. Floxed stop mCAT overexpression mice (Jackson Laboratories, Stock no. 030712) were also bred to generate mCAT; β-LonP1^KO^ mice. *Ins1*^Cre^ alone and mCAT; *Ins1*^Cre^ littermates were utilized as controls for these studies.

### Human islet samples

All human samples were procured from de-identified donors with or without T2D from the Integrated Islet Distribution Program (IIDP) or Prodo Laboratories and studies were approved by the University of Michigan Institutional Review Board. Human primary islets were cultured at 37°C with 5% CO_2_ in PIM(S) media (Prodo Laboratories) supplemented with 10% FBS (Gemini Bio), 100 U/mL penicillin/streptomycin (Gibco), 100 U/mL antibiotic/antimycotic (Gibco), and 1mM PIM(G) (Prodo Laboratories). Islets were used from male and female donors, and donor information is provided in Table S1. Drug treatments used for human islet cultures included 1 μM CDDO (Cayman Chemical), 1 μg/mL tunicamycin (Cayman Chemical), or 5 mM N-acetylcysteine (NAC) for 24 h, respectively, as well as respective vehicle controls (DMSO or PBS).

### Mouse primary islet isolation and culture

Mouse primary islets were isolated by perfusing pancreata with a 1 mg/mL solution of Collagenase P (Millipore Sigma; St Louis, MO, USA) in 1 X HBSS into the pancreatic duct. Following excision of the pancreas, pancreata were incubated at 37°C for 13 min, and Collagenase P was deactivated by addition of 1X HBSS + 10% adult bovine serum (Quench buffer). Pancreata were dissociated mechanically by vigorous shaking for 30 sec, the resulting cell suspension was passed through a 70 μM cell strainer (Fisher Scientific, Waltham, MA, USA). Cells were centrifuged at 1000 rpm for 2 min, the pellet was resuspended in 20 mL Quench buffer and gently vortexed to thoroughly mix. Cells were again centrifuged at 1000 rpm, 1 min. This wash step was repeated once more. Following washes, the cell pellet was resuspended in 5 mL Histopaque (Millipore-Sigma; St Louis, MO, USA) with gentle vortexing. An additional 5 mL Histopaque was layered on the cell suspension, and finally 10 mL Quench buffer was gently layered on top. The cells were spun at 900 x *g* for 30 min at 10°C, with the brake off. The entire Histopaque gradient was pipetted off and passed through an inverted 70 μM filter to trap the islets cells. Islets were washed twice with 10 mL Quench buffer and once with complete islet media (RPMI-1640 supplemented with 100 U/mL penicillin/streptomycin, 10% FBS, 1 mM HEPES, 2 mM L-Glutamine, 100 U/mL antibiotic/antimycotic and 10 mM sodium pyruvate). The filter was inverted into a sterile petri dish and cells were washed into the dish with 4.5 mL complete islet media.

### Pseudoislets

Pseudoislets were prepared using the protocol as described ^73^. Briefly, islets from 3-4-week-old control and β-LonP1^KO^ mice were handpicked and then dispersed with trypsin (Thermo Scientific). Islet cells were counted and incubated with adenovirus expressing an empty vector control (Ad-RIP2 EV) or a protease-deficient LONP1 S855A mutant under control of the rat insulin 2 promoter (Ad-RIP2 S855A) for 2 hr at MOI of 250. Cells were then seeded at 2000 cells per well in CellCarrier Spheroid Ultra-low attachment microplates (PerkinElmer) in enriched pseudoislet media as described ^73^. Cells were allowed to reaggregate for 7 days before being harvested for downstream studies. For *in vitro* LonP1 deletion in adult mouse pseudoislets, islets from 7-8-week-old MIP-Cre^ERT^; *LonP1* ^loxP/loxP^ mice were dispersed and transduced with (Ad-RIP2 EV) and (Ad-RIP2 S855A) for 2 h at MOI of 250, and then seeded at 2000 cells per well in enriched pseudoislet media supplemented with 2 μM 4-hydroxytamoxifen (4-OHT; Sigma) or vehicle control (0.1% ethanol) to induce recombination, followed by biochemical assays. For generation of β-cell enriched human pseudoislets, primary human islets were first dissociated with TrypLE (Gibco) prior to incubation with a mouse monoclonal anti-human NTPDase3 antibody (Ectopeptidases) for 30 min at 4°C. Dissociated human islets were then washed in a magnetic-activated cell sorting buffer (MB) containing 1X sterile phosphate-buffered saline (PBS; Corning), 0.5% FBS (Gemini), and 1% Penicillin-streptomycin solution (Gibco) and then pelleted prior to resuspension in MB containing anti-mouse IgG2a+b MicroBeads (Miltenyi Biotec) and incubation for 15 min at 4°C. NTPDase3+ β-cells were then purified using LS columns (Miltenyi Biotec), followed by counting and incubation with with (Ad-RIP2 EV) and (Ad-RIP2 S855A) for 2 h at MOI of 50, and then seeded at 2000 cells per well in enriched pseudoislet media as above. Cells were allowed to reaggregate for 5 days before incubation with media containing 25 mM glucose + 0.4 mM palmitate (conjugated to BSA as previously described ^97^) or 5mM glucose + BSA alone for 48-72 h prior to downstream studies. Drug treatments used for pseudoislet studies included 40 μM 84-B10 (MedChem Express) for 48 h or 1 μM MKT077 (Cayman Chemical) for 24 h, respectively, as well as respective vehicle controls (DMSO).

### Intraperitoneal Glucose Tolerance Tests (IPGTT)

Mice were fasted for 6 h. Fasting blood glucose measurements were taken by tail nick (Bayer Contour glucometer) before an IP injection of 1 g/kg glucose was administered. Blood glucose measurements were then taken at 15, 30, 60 and 120 mins. Following the test, mice were returned to housing cages with *ad libitum* access to food.

### *In vivo* Glucose Stimulated Insulin Release

Mice were fasted for 6 h. Fasting blood glucose was measured after tail nick with a glucometer (Bayer Contour) and a 20 μL blood sample was collected using capillary tubes (Fisher Scientific) and stored on ice. Mice were injected with 3 g/kg glucose and blood glucose and blood samples were taken after 3 min. Blood samples were ejected from the capillary tubes into 1.5 mL tubes, spun at 16,000 x *g*, 4°C for 10 min, and serum was aliquoted to new 1.5 mL tubes. Serum insulin levels were measured by ELISA (Alpco; Salem, NH). Following the test mice were returned to housing cages with *ad libitum* access to food.

### Intraperitoneal Insulin Tolerance Tests (ITT)

Mice were fasted for 6 h. Fasting blood glucose was measured after tail nick with a glucometer (Bayer Contour). Mice were injected with 0.8 U/kg insulin (Humulin R; Eli Lilly; Indianapolis, IN) and blood glucose measured at 15, 30, and 60 min. Following the test mice were returned to housing cages with *ad libitum* access to food.

### Islet respirometry

Islet respirometry was performed using the BaroFuse instrument (EnTox Sciences, Inc) as previously described.^98^ Setup of the BaroFuse perifusion instrument involved 4 steps: 1. Preheating the system following by filling the Media Reservoirs with pre-equilibrated Media (1X KRBH with 3 mM glucose); 2. Insertion of the Tissue Chamber Assemblies into the Perifusion Module and securing it in place on top of the Media Reservoir to create a gas tight seal; 3. Purging the headspace of the Media Reservoirs with the desired gas composition (typically 5% CO2/21% O2 balance N2) and allow the gas in the headspace to equilibrate with the perifusate in the reservoir; and 4. close the purge port and allow the flow to fill up the Tissue Chambers. When the Tissue Chambers had almost filled, islets were loaded into 6 of the 8 chambers and 2 were left empty to be used as an inflow reference. 200 islets were transferred into the Tissue Chamber (ID = 1.5 mm) with a P-200 pipet. Once islets were loaded, the magnetic stirrers, O2 detector and flow rate monitor are started, and the system and the islets are allowed to equilibrate and establish a stable baseline for 90-120 minutes. Subsequently, test compounds (final concentrations of 20 mM glucose, 10 μM oligomycin, and 3 mM KCN) were injected at precise times. Data are presented as fractional change from baseline (at 3 mM glucose containing media) oxygen concentration rates (OCR) per sample relative to minimum OCR per sample determined following exposure to 3 mM KCN for baseline normalization using the BaroFuse Data Processing Package software (Entox Sciences, Inc.).

### Transmission electron microscopy

Mouse islets were fixed in 2% paraformaldehyde and 2.5% glutaraldehyde in 0.1 M sodium cacodylate (pH 7.4) for 5 minutes at room temperature and then stored at 4°C for further processing. Following agarose embedding, islets were then subjected to osmification in 1.5% K_4_Fe(CN)_6_ + 2% OsO_4_ in 0.1 cacodylate buffer for 1 h, dehydrated by serial diluted ethanol solutions (30, 50, 70, 80, 90, 95 and 100%) and embedded in Spurr’s resin by polymerization at 60°C for 24 h. Polymerized resins were then sectioned at 90 nm thickness using a Leica EM UC7 ultramicrotome and imaged at 70 kV by using a JEOL 1400 transmission electron microscopy equipped with an AMT CMOS imaging system. Mitochondrial morphology was analyzed and quantified by Fiji, as previously described.^17^

### Biochemical assessments of protein solubility

Mouse islets were lysed with Triton X-100 buffer (20 mM Tris-HCl, pH 7.4, 150 mM NaCl, 2 mM EDTA, 1% Triton X-100) supplemented with protease and phosphatase inhibitors (Protease inhibitor set III and Phosphatase inhibitor set II, Calbiochem), incubated on ice for 30 min, and then centrifuged at 21,000 g for 10 min at 4°C to separate into a supernatant (Triton X-100 soluble fraction) and a pellet (Triton X-100 insoluble fraction). Insoluble fractions were dissolved in 2% SDS containing a 50% volume of Triton X-100 buffer (with protease/phosphatase inhibitors), followed by immunoblotting or TMT labeling mass spectrometry.

### SDS-PAGE

Isolated mouse islets or human pseudoislets were homogenized in RIPA buffer containing protease and phosphatase inhibitors. Samples were centrifuged at full speed for 10min, 4 °C to pellet insoluble material and the supernatant was used for immunoblot analysis. Protein quantification was carried out using a Pierce MicroBCA kit (ThermoFisher). Protein lysates were prepared in Laemmli buffer with DTT and denatured at 70°C for 10min, or 37°C for 30min for OXPHOS analysis. Samples were then run on a 4-15% Tris-glycine protein gel (Bio-Rad) at 150V until separated. Samples were then transferred to a nitrocellulose membrane at 90V for 90min, membranes were blocked with 5% skim milk in 1X TBS + 0.05% Tween-20 for 1 h, and incubated overnight at 4 °C with primary antisera, including LONP1 (1:1000, ProteinTech, Catalog# 15440-1-AP), GRP75 (mtHSP70, 1:1000, ProteinTech, Catalog# 14887-1-AP), HSP60 (1:1000, Santa Cruz, sc-13115), DNAJA3 (1:1000, Santa Cruz, sc-18820), CLPP (1:1000, ProteinTech, 15698-1-AP), Aconitase 2 (1:1000, Abcam, ab110321), Anti-NDUFA10 (1:1000, Santa Cruz, sc376357), SDHA (1:1000, Abcam, ab14715), Total OXPHOS rodent antibody cocktail (1:1000; Abcam, ab110413), Total OXPHOS human antibody cocktail (1:1000; Abcam, ab110411), TOM20 (1:1000; Cell Signaling Technology, Catalog# 42406), VDAC1 (1:1000; Abcam, ab14734), Catalase (1:2000, Athens Research &Technology, Catalog# 01-05-30000), phospho-γH2AX (1:1000, Millipore, Catalog# 05-636), VINCULIN (1:1000; Millipore, Catalog# CP74). Membranes were then washed 3X with 1X TBS+0.05% Tween-20 and incubated with species-specific HRP conjugated secondary antisera (Vector Labs).

### Blue-native PAGE

Blue-native PAGE was performed similar to a previously published protocol as described.^71^ Briefly, mouse islets were lysed in native sample buffer (62.5 mM Tris-HCl, 10% glycerol and protease inhibitors, pH 7.0) containing 1% n-Dodecyl-B-D-Maltoside (DDM; Sigma and 1% digitonin (Sigma), incubated on ice for 30 min, and centrifuged at 21,000 g at 4℃ for 15 min. Equal quantites of protein lysates, determined by microBCA assay kit, (Pierce) were added to 4-15% polyacrylamide gradient gels and supplemented with Coomassie Brilliant blue (0.2% final concentration; ThermoFisher) added to the supernatant. Following electrophoresis, proteins were transferred to activated PVDF membranes, followed by immunoblotting. Immunoblotting was performed by using Anti-NDUFA9 (1:1000, Abcam, ab14713), SDHA (1:1000, Abcam, ab14715), Anti-UQCRC2 (1:1000, Abcam, ab14745), Total OXPHOS blue native WB antibody cocktail (1:1000; Abcam, ab110412), and species-specific HRP-conjugated secondary antibodies (Vector Laboratories).

### β-cell mass analysis

Whole mouse pancreas was excised, weighed, and fixed in 4% paraformaldehyde for 16 h at 4°C. Samples were stored in 70% ethanol, 4°C before being embedded in paraffin and sectioned. 3 independent depths of sections, at least 50 μM apart, were dewaxed, and rehydrated and antigen retrieval was carried out using 10 mM sodium citrate (pH 6.0) in a microwave for 10 min. Sections were washed twice with 1X PBS, blocked for 1 h at room temperature with 5% donkey serum in PBT (1X PBS, 0.1% Triton X-100, 1% BSA). Sections were then incubated in the following primary antisera overnight at 4°C in PBT: guinea pig anti-insulin (Thermo; PA1-26938. Sections were then washed twice with PBS and incubated for 2 h at room temperature with species specific Cy2 and Cy3 conjugated secondary antibodies. Nuclear labelling was performed using DAPI (Molecular Probes). Sections were scanned using an Olympus IX81 microscope (Olympus; Center Valley, PA, USA) at 10X magnification, with image stitching for quantification (Olympus). β-cell mass quantification (estimated as total insulin positive area/total pancreatic area multiplied by pancreatic weight) was performed on stitched images of complete pancreatic sections from 3 independent regions.

### Immunofluorescence imaging

Whole mouse pancreas was excised, weighed, and fixed in 4% paraformaldehyde for 16 h at 4°C. Samples were stored in 70% ethanol at 4°C, before being embedded in paraffin and sectioned. Following dewaxing and rehydration, antigen retrieval was performed in 10 mM citric acid (pH 6.0) or HistoVT buffer (Nacalai) as previously described.^51^ Samples were then blocked in PBT containing 10% donkey serum or Blocking One solution (Nacalai) for 1 h at room temperature. Pancreatic sections were stained with the following primary antibodies, insulin (Thermo-Fisher, PA1-26938 or Fitzgerald, 20-IP35), Ki67 (Abcam, ab16667), PDX1 (Abcam, ab47383), ALDH1A3 (Novus Bio, NBP2-15339), Urocortin 3 (Salk PBL7218, a gift from the laboratory of Dr. Wylie Vale), GRP75 (mtHSP70, ProteinTech, 14887-1-AP), SDHA (Abcam, ab14715), as well as species specific Cy2 and Cy3 conjugated secondary antibodies (Jackson Immunoresearch). Nuclear labelling was performed using DAPI (Molecular Probes). High magnification images were captured under oil immersion on an Olympus IX81 microscope.

### Imaging of mitochondrial morphology in mouse β-cells

Images of pancreas sections stained with SDHA and Insulin (to identify β-cells) were captured with an IX81 microscope (Olympus) using an ORCA Flash4 CMOS digital camera (Hamamatsu). Immunostained pancreatic sections, were captured with Z-stack images and subjected to deconvolution (CellSens; Olympus). Quantitative 3D assessments of mitochondrial morphology and network were performed on ImageJ using Mitochondria Analyzer plugin ^99^. Co-localization analyses were performed on Z-stack images of immunostained dissociated islet cells using the Coloc2 plugin on ImageJ.

### Quantitative PCR of cDNA and mtDNA content

DNA and RNA samples were isolated from mouse islets with the Blood/Tissue DNeasy kit (Qiagen) and MicroElute total RNA kit, respectively. DNase-treated RNA (Ambion DNA free kit) was reverse transcribed using High-Capacity cDNA Reverse Transcription Kit (Thermo Fisher Scientific) as previously described ^100^. Quantitative PCR (qPCR) was performed with SYBR-based detection (Universal SYBR Green Supermix; Bio-Rad) using primers for mt9/11 for mtDNA (5′- GAGCATCTTATCCACGCTTCC-3′ and 5′- GGTGGTACTCCCGCTGTAAA-3′) and *Ndufv1* (5′- CTTCCCCACTGGCCTCAAG-3′ and 5′- CCAAAACCCAGTGATCCAGC-3′) for nuclear DNA; For gene expression analysis of cDNA, primers included *CHOP* (5′- CCTAGCTTGGCTGACAGAGG-3′ and 5′-CTGCTCCTTCTCCTTCATGC-3′), *GRP75* (5′- AGGGCAAACAAGCAAAGGTCC-3′ and 5′-TGGTGACAGCTTGCCGTTTTG-3′), *DNAJA3* (5′- AGAACCATGGATAGCTCCGCA-3′ and 5′-TCCAGTTGACCGCTTTCCTCA-3′), *HSP60* (5′- ACAATGGGGCCAAAGGGAAGA-3′ and 5′-GACTTTGCAACAGTGACCCCA-3′), *HSP10* (5′- GACTTTGCAACAGTGACCCCA-3′ and 5′- CTTTGGTGCCTCCATATTCTGGG-3′), *CLPP* (5′- GAACTGCGACGCGAGCTTTC-3′ and 5′- CACACTGTCGTCAATCGGGC-3′), and *HPRT* (5’- GGCCAGACTTTGTTGGATTTG-3′ and 5′- TGCGCTCATCTTAGGCTTTGT-3′).

### TMT labeling mass spectrometry (MS)-based quantitative proteomics

Human islets from donors with or without type 2 diabetes were collected and lysed with Triton X-100 buffer containing protease and phosphatase inhibitors as described above for TMT labeling mass spectrometry.

#### Protein Digestion and TMT labeling

Depending on the solubilization buffer, proteins from insoluble fractions were initially processed via filter-aided sample preparation (FASP) protocol using FASP protein digestion kit (Abcam; #ab270519)) and proteins from soluble fractions were processed via standard TMT sample processing protocol (Thermo-Fisher). Protein samples were digested with trypsin/Lys-C mix (1:25 protease: protein; Promega) at 37°C was performed with constant mixing using a thermomixer. The TMT 16-plex reagents were dissolved in 20 μl of anhydrous acetonitrile and labeling was performed by transferring the entire digest to TMT reagent vial and incubating at room temperature for 1 h. Reaction was quenched by adding 5 μl of 5% hydroxyl amine and further 15 min incubation. Labeled samples were mixed and dried using a vacufuge. An offline fractionation of the combined sample (∼200 μg) into 8 fractions was performed using high pH reversed-phase peptide fractionation kit according to the manufacturer’s protocol (Pierce; Cat #84868). Fractions were dried and reconstituted in 9 μl of 0.1% formic acid / 2% acetonitrile in preparation for LC-MS/MS analysis.

#### Liquid chromatography-mass spectrometry analysis (LC-multinotch MS3)

To obtain superior quantitation accuracy, we employed multinotch-MS3 (McAlister GC, reference below) which minimizes the reporter ion ratio distortion resulting from fragmentation of co-isolated peptides during MS analysis. Orbitrap Fusion (Thermo-Fisher) and RSLC Ultimate 3000 nano-UPLC (Dionex) was used to acquire the data. 2 μl of the sample was resolved on a PepMap RSLC C18 column (75 μm i.d. x 50 cm; Thermo-Fisher) at the flowrate of 300 nl/min using 0.1% formic acid/acetonitrile gradient system (2-22% acetonitrile in 150 min; 22-32% acetonitrile in 40 min; 20 min wash at 90% followed by 50 min re-equilibration) and directly spray onto the mass spectrometer using EasySpray source (Thermo-Fisher). Mass spectrometer was set to collect one MS1 scan (Orbitrap; 120K resolution; AGC target 2 x 10^5^; max IT 100 ms) followed by data-dependent, “Top Speed” (3 s) MS2 scans (collision induced dissociation; ion trap; NCE 35; AGC 5 x 10^3^; max IT 100 ms). For multinotch-MS3, top 10 precursors from each MS2 were fragmented by HCD followed by Orbitrap analysis (NCE 55; 60K resolution; AGC 5 x 10^4^; max IT 120 ms, 100-500 m/z scan range).

#### Data analysis

Proteome Discoverer (v2.4; Thermo Fisher) was used for data analysis. MS2 spectra were searched against SwissProt human protein database (20291 entries; reviewed; downloaded on 12/13/2021) using the following search parameters: MS1 and MS2 tolerance were set to 10 ppm and 0.6 Da, respectively; carbamidomethylation of cysteines (57.02146 Da) and TMT labeling of lysine and N-termini of peptides (304.2071 Da) were considered static modifications; oxidation of methionine (15.9949 Da) and deamidation of asparagine and glutamine (0.98401 Da) were considered variable. Identified proteins and peptides were filtered to retain only those that passed ≤ 1% FDR threshold. Quantitation was performed using high-quality MS3 spectra (Average signal-to-noise ratio of 16 and < 50% isolation interference).

#### TMT-MS data analysis

TMT-Integrator was applied for additional identification assessment and data processing to achieve accurate quantification. The quantification reports generated by TMT-Integrator were used for downstream analysis. Differential expression (DE) analysis was conducted for all identified proteins under different treatments compared to DMSO, using the linear model provided by the limma package. Functional analysis of Gene Ontology (GO) terms among all proteins was performed using the Cluster Profiler package. To visualize the DE results, volcano plots were generated using the R package ggplot2. The log2 fold change (log2FC) was represented by Limma’s moderated t-statistic on the x-axis, and the -log10 transformed p-value (-log10 p-value) on the y-axis. Mitocarta 3.0 targets were highlighted by green color points.

### Adenoviral vectors and viral preparation

FLAG-tagged human LONP1 S855A was generated by PCR amplification of pQCXIP-LONP1 S855A (kindly provided by David Chan, California Institute of Technology) utilizing PCR primers to add a FLAG epitope tag in frame at the C-terminus of LONP1. The PCR amplified fragment was subcloned into pCR-Blunt II-TOPO (Zero blunt TOPO cloning kit, Invitrogen) for sequence validation by Sanger sequencing (Eurofins), followed by restriction digestion for KpnI and MfeI, and T4 ligation of fragments into a linearized pAd-RIP2 vector (kindly provided by Dr. Leslie Satin, University of Michigan) for further adenovirus generation processing in Vector Core at University of Michigan. Briefly, pAd-RIP2-LONP1 S855A or pAd-RIP2 empty vector plasmids were co-transfected and recombined with cAd5 (E1-E3 genes deleted). Individual plaques were selected, and viral lysates were screened for protein expression. Selected clones were then expanded in 911 cells and purified using CsCl gradient ultracentrifugation. Viral bands were collected and flushed through a Sepharose CL-4B (Cytiva, Sweden) column. Eluted adenovirus was diluted in a sterile isosaline solution (10 mM Tris-HCl, 137 mM NaCl, 5 mM KCl, 1 mM MgCl_2_ and 10% glycerol (v/v)). Titers were determined by counting plaque forming units from 911 cells infected with serially diluted adenovirus and overlayed with 0.8% Noble Agar in 1x EMEM + 5% FBS.

### Cell death and ROS assessments

Cytoplasmic histone-complexed DNA fragments were determined in islets using the Cell Death ELISAplus (Roche), per the manufacturer’s protocol. Cell pellets were analyzed to detect apoptosis. ELISA absorbance was measured at 450nm and normalized to total DNA content (SYBRGreen; Invitrogen) using an absorbance/fluorescence microplate reader (BioTek). Terminal deoxy-nucleotidyl transferase dUTP nick end labeling (TUNEL) was performed on transduced human islets (Apoptag In situ Apoptosis Detection Kit; Millipore) as per the manufacturer’s protocol. Human islets were prepared for staining as previously described ^97^. TUNEL stained samples were counter-stained with insulin antibodies. Images were captured with an IX81 microscope (Olympus) using an ORCA Flash4 CMOS digital camera (Hamamatsu). ROS levels were measured by a cell-based fluorogenic assay kit (ROS/Superoxide detection assay kit; Abcam) as per the manufacturer’s protocol as previously described ^95^.

### Statistics

All data are presented as mean ± SEM. Statistical analyses were performed in GraphPad Prism 10.0 software by using two-tailed Student’s *t* tests, one-way or two-way ANOVA, followed by Tukey’s or Sidak’s post-hoc test for multiple comparisons as appropriate. Outlier tests (ROUT method ^101^) were routinely performed in GraphPad Prism. A P value < 0.05 was considered significant.

## RESOURCE AVAILABILITY

### Lead Contact

Further information and request for resources and reagents should be directed to and will be fulfilled by the lead contact, Scott Soleimanpour

### Materials Availability

LonP1^loxP^ mice are available on request from the lead author.

### Data Availability

Proteomics data deposition at ProteomeXchange is in process and will be publicly available at the date of publication. Microscopy data will be shared by lead contact on request.

Any additional information required to reanalyze the data reported in this paper is available from the lead author on request.

## Acknowledgements

S.A.S. acknowledges support from the JDRF (SRA-2023-1392), the NIH (R01 DK108921, R01 DK135032, R01 DK135268, R01 DK136671, R01 DK127270, U01 DK127747, P30 DK020572), the Department of Veterans Affairs (I01 BX004444), the Brehm family, and the Anthony family. E.M.W. was supported by the NIH (5K01DK133533). M.L.S. acknowledges support from the ADA Pathway to Stop Diabetes Accelerator Award (1-81-ACE-015) and the NIH (R01DK136671, R01DK118011). The JDRF Career Development Award to S.A.S. is partly supported by the Danish Diabetes Academy and the Novo Nordisk Foundation. We acknowledge the Microscopy, Imaging and Cellular Physiology Core and Islet Core of the University of Michigan DRC (P30 DK020572) for assistance with imaging studies and pseudoislet studies, respectively. Proteomics studies were carried out in the Proteomics Resource Facility at the University of Michigan. Human pancreatic islets and/or other resources were provided by the NIDDK-funded Integrated Islet Distribution Program (IIDP) (RRID:SCR _014387) at City of Hope, NIH Grant # U24DK098085. We thank Dr. K. Claiborn and members of the Soleimanpour laboratory for helpful advice.

## Author Contributions

J.L. conceived, designed, and performed experiments, interpreted results, drafted and reviewed the manuscript. J.Z., designed and performed experiments, and interpreted results. Y.D. interpreted results and reviewed the manuscript. E.C.R. designed and performed experiments and interpreted results. E.M.W. and V.S. designed and performed experiments, interpreted results and reviewed the manuscript. D.H. performed experiments, interpreted results, and reviewed the manuscript. M.B.P, K.B., E.M., S.N., and R.K. designed and performed experiments, and interpreted results. V.K. designed and performed experiments, interpreted results and reviewed the manuscript. A.I.N, M.L.S., and D.C.C designed studies, interpreted results, edited, and reviewed the manuscript. S.A.S. conceived and designed the studies, interpreted results, drafted, edited, and reviewed the manuscript.

