## Supplementary Data for "LONP1 regulation of mitochondrial protein folding provides insight into beta cell failure in type 2 diabetes"

**Table S1.** Clinical characteristics of human islet donors

| Unique Islet Prep Identifier | Islet Isolation Center | Age (year) | BMI | Sex | T2D | HbA1c (%) | Cause of Death |
|---|---|---|---|---|---|---|---|
| HP-21181-01T2D | Prodo | 47 | 31.2 | M | Yes | 6.6 | Stroke |
| RRID: SAMN20567145 | IIDP | 58 | 22.9 | M | Yes | 10.5 | Cerebrovascular/stroke |
| HP-21263-01T2D | Prodo | 62 | 34.6 | M | Yes | 6.8 | Stroke |
| HP-21342-01T2D | Prodo | 48 | 39.2 | M | Yes | 7.1 | Anoxia |
| RRID: SAMN23079315 | IIDP | 37 | 30.2 | F | No | NA | Anoxia |
| RRID: SAMN23245917 | IIDP | 54 | 26.8 | F | No | NA | Cerebrovascular/stroke |
| HP-21324-01 | Prodo | 44 | 25.7 | M | No | 5.8 | Trauma |
| HP-21364-01 | Prodo | 55 | 24.9 | M | No | 5.4 | Stroke |
| HP-22362-01 | Prodo | 51 | 24.9 | M | No | 5.3 | Stroke |
| HP-23019/20-01 | Prodo | 55 | 30.4 | M | No | 5.6 | Anoxia |
| HP-23312-01 | Prodo | 37 | 29.6 | F | No | 5.1 | Anoxia |
| HP-23350-01 | Prodo | 71 | 29.3 | F | No | 5.1 | Stroke |
| HP-23301-01 | Prodo | 64 | 25.5 | M | No | 5.2 | Stroke |
| HP-23305-01 | Prodo | 47 | 28.3 | M | No | 5.3 | Stroke |
| RRID: SAMN38227365 | IIDP | 61 | 24.2 | M | No | 5.4 | Cerebrovascular/stroke |
| HP-23342-01 | Prodo | 34 | 24.6 | M | No | 5.4 | Anoxia |

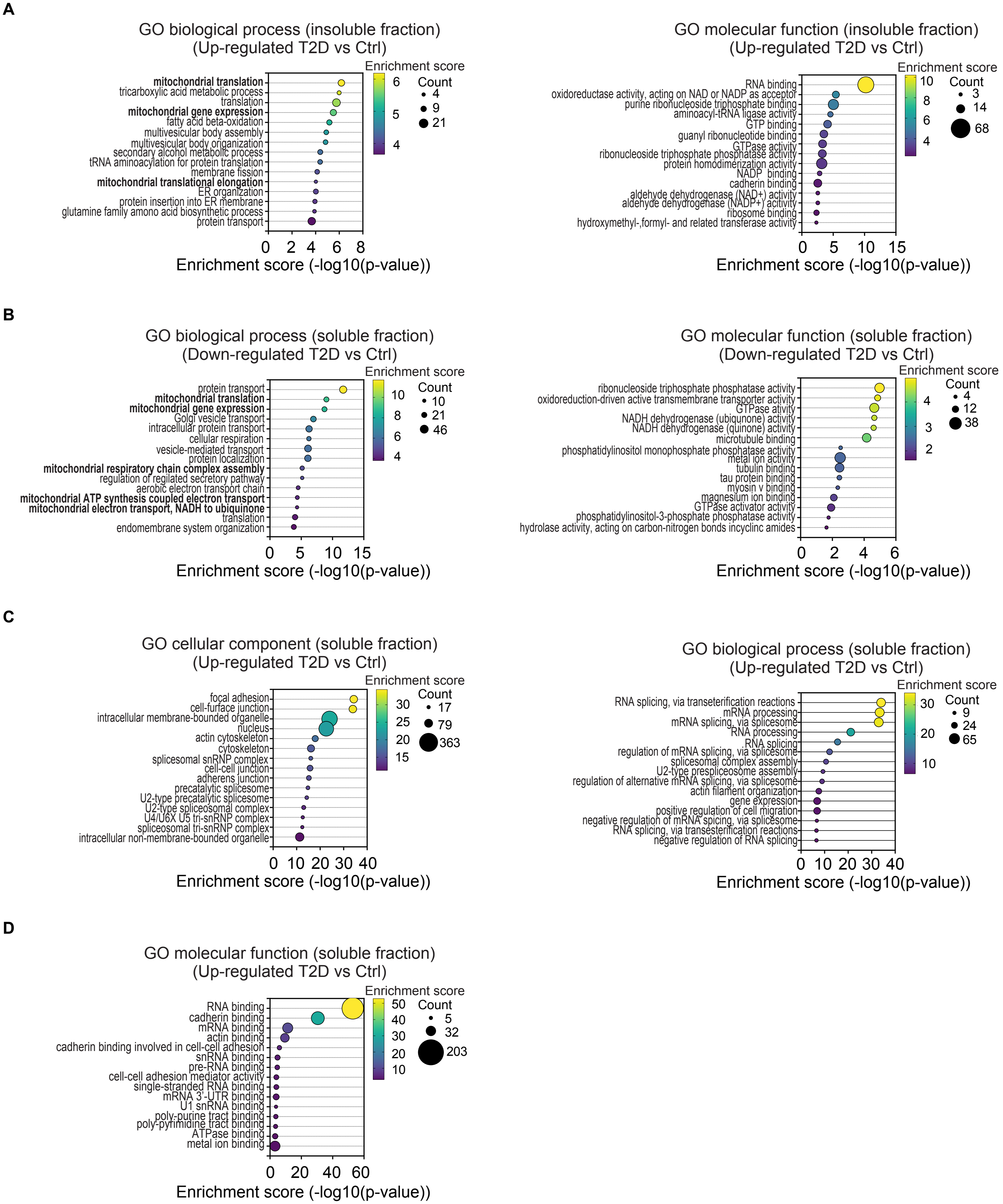
A
GO biological process (insoluble fraction)
(Up-regulated T2D vs Ctrl)
mitochondrial translation
tricarboxylic acid metabolic process
translation
mitochondrial gene expression
fatty acid beta-oxidation
multivesicular body assembly
multivesicular body organization
secondary alcohol metabolic process
tRNA aminoacylation for protein translation
membrane fission
mitochondrial translational elongation
ER organization
protein insertion into ER membrane
glutamine family amono acid biosynthetic process
protein transport
Enrichment score
Count
Enrichment score (-log10(p-value))
GO molecular function (insoluble fraction)
(Up-regulated T2D vs Ctrl)
RNA binding
oxidoreductase activity, acting on NAD or NADP as acceptor
purine ribonucleoside triphosphate binding
aminoacyl-tRNA ligase activity
GTP binding
guanyl ribonucleotide binding
GTPase activity
ribonucleoside triphosphate phosphatase activity
protein homodimerization activity
NADP binding
cadherin binding
aldehyde dehydrogenase (NAD+) activity
aldehyde dehydrogenase (NADP+) activity
ribosome binding
hydroxymethyl-,formyl- and related transferase activity
B
GO biological process (soluble fraction)
(Down-regulated T2D vs Ctrl)
protein transport
mitochondrial translation
mitochondrial gene expression
Golgi vesicle transport
intracellular protein transport
cellular respiration
vesicle-mediated transport
protein localization
mitochondrial respiratory chain complex assembly
regulation of regilated secretory pathway
aerobic electron transport chain
mitochondrial ATP synthesis coupled electron transport
mitochondrial electron transport, NADH to ubiquinone
translation
endomembrane system organization
GO molecular function (soluble fraction)
(Down-regulated T2D vs Ctrl)
ribonucleoside triphosphate phosphatase activity
oxidoreduction-driven active transmembrane transporter activity
GTPase ativity
NADH dehydrogenase (ubiquinone) activity
NADH dehydrogenase (quinone) activity
microtubule binding
phosphatidylinositol monophosphate phosphatase activity
metal ion activity
tubulin binding
tau protein binding
myosin v binding
magnesium ion binding
GTPase activator activity
phosphatidylinositol-3-phosphate phosphatase activity
hydrolase activity, acting on carbon-nitrogen bonds incyclinc amides
C
GO cellular component (soluble fraction)
(Up-regulated T2D vs Ctrl)
focal adhesion
cell-furface junction
intracellular membrane-bounded organelle
nucleus
actin cytoskeleton
cytoskeleton
spliceosomal snRNP complex
cell-cell junction
adherens junction
precatalytic spliceosome
U2-type precatalytic spliceosome
U2-type spliceosomal complex
U4/U6X U5 tri-snRNP complex
spliceosomal tri-snRNP complex
intracellular non-membrane-bounded organelle
GO biological process (soluble fraction)
(Up-regulated T2D vs Ctrl)
RNA splicing, via transeterification reactions
mRNA processing
mRNA splicing, via spliceosome
RNA processing
RNA splicing
regulation of mRNA splicing, via spliceosome
spliceosomal complex assembly
U2-type prespliceosome assembly
regulation of alternative mRNA splicing, via spliceosome
actin filament organization
gene expression
positive regulation of cell migration
negative regulation of mRNA spicing, via spliceosome
RNA splicing, via transesterification reactions
negative regulation of RNA splicing
D
GO molecular function (soluble fraction)
(Up-regulated T2D vs Ctrl)
RNA binding
cadherin binding
mRNA binding
actin binding
cadherin binding involved in cell-cell adhesion
snRNA binding
pre-RNA binding
cell-cell adhesion mediator activity
single-stranded RNA binding
mRNA 3'-UTR binding
U1 snRNA binding
poly-purine tract binding
poly-pyrimidine tract binding
ATPase binding
metal ion binding

**Figure S1. Pathway analyses of quantitative proteomics of soluble and insoluble fractions from human islet donors with or without T2D.**

(A) GO biological process (left) and molecular function (right) analysis of significantly upregulated insoluble proteins from human islet donors with T2D compared to non-diabetic controls. n = 4 independent islet donors/group. (B**)** GO biological process (left) and molecular function (right) analysis of significantly downregulated insoluble proteins from human islet donors with T2D compared to non-diabetic controls. n = 4 independent islet donors/group. (C) GO cellular component (left) and biological process (right) analysis of significantly upregulated soluble proteins from human islet donors with T2D compared to non-diabetic controls. n = 4 independent islet donors/group. (D) GO molecular function analysis of significantly upregulated soluble proteins from human islet donors with T2D compared to non-diabetic controls. n = 4 independent islet donors/group.

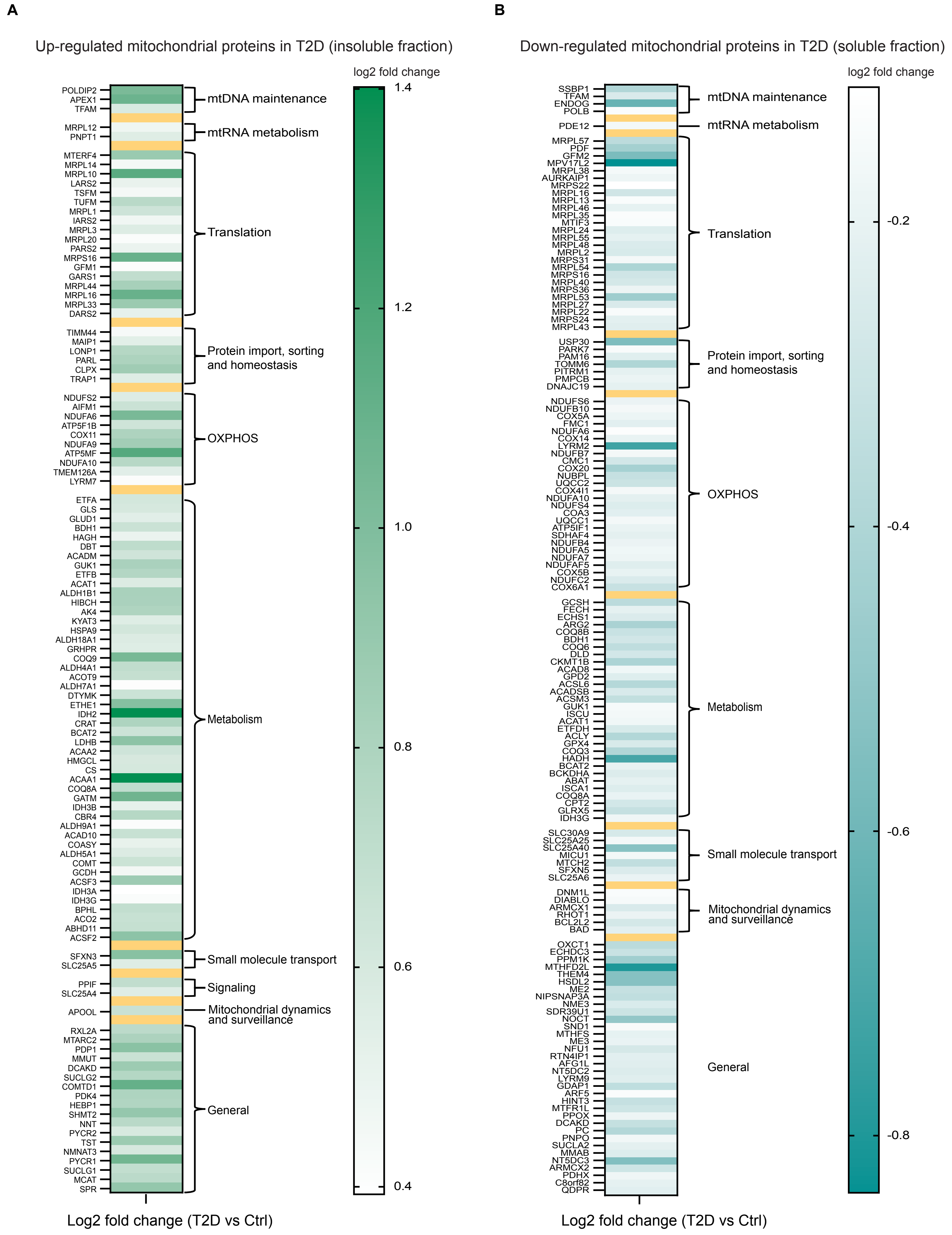

A
Up-regulated mitochondrial proteins in T2D (insoluble fraction)
log2 fold change
1.4
1.2
1.0
0.8
0.6
0.4
POLDIP2
APEX1
TFAM
mtDNA maintenance
MRPL12
PNPT1
mtRNA metabolism
MTERF4
MRPL14
MRPL10
LARS2
TSFM
TUFM
MRPL1
IARS2
MRPL3
MRPL20
PARS2
MRPS16
GFM1
GARS1
MRPL44
MRPL16
MRPL33
DARS2
Translation
TIMM44
MAIP1
LONP1
PARL
CLPX
TRAP1
Protein import, sorting and homeostasis
NDUFS2
AIFM1
NDUFA6
ATP5F1B
COX11
NDUFA9
ATP5MF
NDUFA10
TMEM126A
LYRM7
OXPHOS
ETFA
GLS
GLUD1
BDH1
HAGH
DBT
ACADM
GUK1
ETFB
ACAT1
ALDH1B1
HIBCH
AK4
KYAT3
HSPA9
ALDH18A1
GRHPR
COQ9
ALDH4A1
ACOT9
ALDH7A1
DTYMK
ETHE1
IDH2
CRAT
BCAT2
LDHB
ACAA2
HMGCL
CS
ACAA1
COQ8A
GATM
IDH3B
CBR4
ALDH9A1
ACAD10
COASY
ALDH5A1
COMT
GCDH
ACSF3
IDH3A
IDH3G
BPHL
ACO2
ABHD11
ACSF2
Metabolism
SFXN3
SLC25A5
Small molecule transport
PPIF
SLC25A4
Signaling
APOOL
Mitochondrial dynamics and surveillance
RXL2A
MTARC2
PDP1
MMUT
DCAKD
SUCLG2
COMTD1
PDK4
HEBP1
SHMT2
NNT
PYCR2
TST
NMNAT3
PYCR1
SUCLG1
MCAT
SPR
General
Log2 fold change (T2D vs Ctrl)
B
Down-regulated mitochondrial proteins in T2D (soluble fraction)
log2 fold change
-0.2
-0.4
-0.6
-0.8
SSBP1
TFAM
ENDOG
POLB
mtDNA maintenance
PDE12
mtRNA metabolism
MRPL57
PDF
GFM2
MPV17L2
MRPL38
AURKAIP1
MRPS22
MRPL16
MRPL13
MRPL46
MRPL35
MTIF3
MRPL24
MRPL55
MRPL48
MRPL2
MRPS31
MRPL54
MRPS16
MRPL40
MRPS36
MRPL53
MRPL27
MRPL22
MRPS24
MRPL43
Translation
USP30
PARK7
PAM16
TOMM6
PITRM1
PMPCB
DNAJC19
Protein import, sorting and homeostasis
NDUFS6
NDUFB10
COX5A
FMC1
NDUFA6
COX14
LYRM2
NDUFB7
CMC1
COX20
NUBPL
UQCC2
COX4I1
NDUFA10
NDUFS4
COA3
UQCC1
ATP5IF1
SDHAF4
NDUFB4
NDUFA5
NDUFA7
NDUFAF5
COX5B
NDUFC2
COX6A1
OXPHOS
GCSH
FECH
ECHS1
ARG2
COQ8B
BDH1
COQ6
DLD
CKMT1B
ACAD8
GPD2
ACSL6
ACADSB
ACSM3
GUK1
ISCU
ACAT1
ETFDH
ACLY
GPX4
COQ3
HADH
BCAT2
BCKDHA
ABAT
ISCA1
COQ8A
CPT2
GLRX5
IDH3G
Metabolism
SLC30A9
SLC25A25
SLC25A40
MICU1
MTCH2
SFXN5
SLC25A6
Small molecule transport
DNM1L
DIABLO
ARMCX1
RHOT1
BCL2L2
BAD
Mitochondrial dynamics and surveillance
OXCT1
ECHDC3
PPM1K
MTHFD2L
THEM4
HSDL2
ME2
NIPSNAP3A
NME3
SDR39U1
NOCT
SND1
MTHFS
ME3
NFU1
RTN4IP1
AFG1L
NT5DC2
LYRM9
GDAP1
ARF5
HINT3
MTFR1L
PPOX
DCAKD
PC
PNPO
SUCLA2
MMAB
NT5DC3
ARMCX2
PDHX
C8orf82
QDPR
General
Log2 fold change (T2D vs Ctrl)

**Figure S2. MitoPathways assessment of soluble and insoluble proteins from islet donors with T2D.**

(A) Differential expression heatmap of significantly upregulated mitochondrial proteins in the insoluble fraction of T2D islets compared to non-diabetic controls. n = 4 independent islet donors/group. Proteins are categorized based on annotation from MitoPathways3.0. (B) Differential expression heatmap of significantly downregulated mitochondrial proteins in the soluble fraction of T2D islets compared to non-diabetic controls. Proteins are categorized based on annotation from MitoPathways3.0.

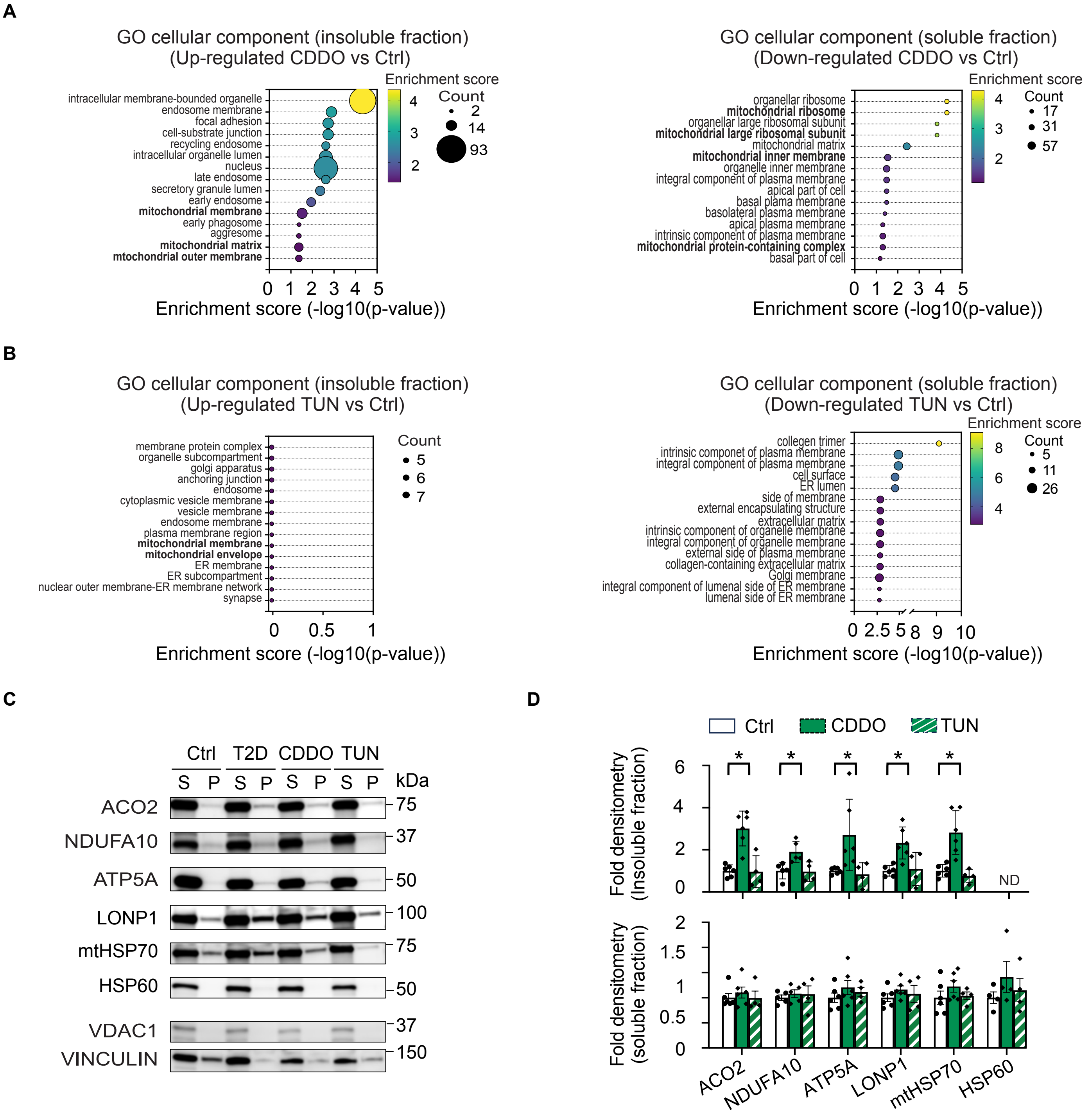

A
GO cellular component (insoluble fraction)
(Up-regulated CDDO vs Ctrl)
Enrichment score
Count
intracellular membrane-bounded organelle
endosome membrane
focal adhesion
cell-substrate junction
recycling endosome
intracellular organelle lumen
nucleus
late endosome
secretory granule lumen
early endosome
mitochondrial membrane
early phagosome
aggresome
mitochondrial matrix
mtochondrial outer membrane
Enrichment score (-log10(p-value))
GO cellular component (soluble fraction)
(Down-regulated CDDO vs Ctrl)
organellar ribosome
mitochondrial ribosome
organellar large ribosomal subunit
mitochondrial large ribosomal subunit
mitochondrial matrix
mitochondrial inner membrane
organelle inner membrane
integral component of plasma membrane
apical part of cell
basal plama membrane
basolateral plasma membrane
apical plasma membrane
intrinsic component of plasma membrane
mitochondrial protein-containing complex
basal part of cell
Enrichment score (-log10(p-value))
B
GO cellular component (insoluble fraction)
(Up-regulated TUN vs Ctrl)
Count
membrane protein complex
organelle subcompartment
golgi apparatus
anchoring junction
endosome
cytoplasmic vesicle membrane
vesicle membrane
endosome membrane
plasma membrane region
mitochondrial membrane
mitochondrial envelope
ER membrane
ER subcompartment
nuclear outer membrane-ER membrane network
synapse
Enrichment score (-log10(p-value))
GO cellular component (soluble fraction)
(Down-regulated TUN vs Ctrl)
Enrichment score
Count
collegen trimer
intrinsic componet of plasma membrane
integral component of plasma membrane
cell surface
ER lumen
side of membrane
external encapsulating structure
extracellular matrix
intrinsic component of organelle membrane
integral component of organelle membrane
external side of plasma membrane
collagen-containing extracellular matrix
Golgi membrane
integral component of lumenal side of ER membrane
lumenal side of ER membrane
Enrichment score (-log10(p-value))
C
Ctrl T2D CDDO TUN
S P S P S P S P
kDa
ACO2 75
NDUFA10 37
ATP5A 50
LONP1 100
mtHSP70 75
HSP60 50
VDAC1 37
VINCULIN 150
D
Ctrl CDDO TUN
Fold densitometry (Insoluble fraction)
ND
Fold densitometry (soluble fraction)
ACO2
NDUFA10
ATP5A
LONP1
mtHSP70
HSP60

**Figure S3. Alterations in mitochondrial protein solubility are observed in human islets following pharmacologic induction of mitochondrial rather than ER protein misfolding.**

(A) Left - cellular component analysis of significantly upregulated insoluble proteins from human islets following exposure to 1 μM CDDO compared to vehicle (DMSO) control islets. Right - cellular component analysis of significantly downregulated soluble proteins from human islets following exposure to 1 μM CDDO compared to vehicle (DMSO) control islets. n = 4 independent islet donors/group. (B) Left - cellular component analysis of significantly upregulated insoluble proteins from human islets following exposure to 1 μg/mL tunicamycin (TUN) compared to vehicle (DMSO) control islets. Right - cellular component analysis of significantly downregulated soluble proteins from human islets following exposure to 1 μg/mL TUN compared to vehicle (DMSO) control islets. n = 4 independent islet donors/group. (C) Expression of selected mitochondrial proteins by WB of soluble and insoluble fractions of human islets isolated from donors with T2D and non-diabetic control donors (Ctrl) as well as human islets from non-diabetic control donor exposed to CDDO or TUN. Representative image of 4 independent human islet donors/group. Expression of select mitochondrial proteins from both soluble and insoluble fractions of T2D islets and non-diabetic control donor islets with CDDO or TUN exposure by WB. Representative images of 4 independent islet donors/group. VDAC1 serves as a soluble mitochondrial protein loading control. VINCULIN serves as a loading control for both soluble and insoluble fractions. S, soluble fraction; P, insoluble fraction. (D) Quantification of protein expression by densitometry (normalized to VINCULIN) from studies in Figure S3C. n = 4 independent islet donors/group. $^{*}p < 0.05$ by one-way ANOVA followed by Tukey's multiple comparisons test. All data in figure are presented as mean ± SEM.

A

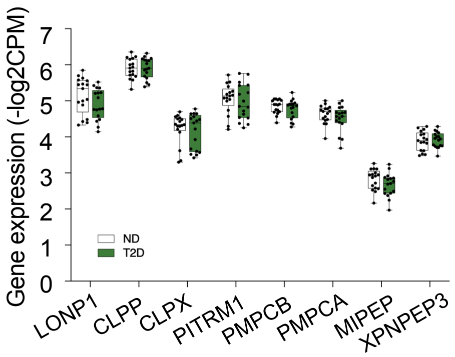
Gene expression (-log2CPM)
7
6
5
4
3
2
1
0
ND
T2D
LONP1
CLPP
CLPX
PITRM1
PMPCB
PMPCA
MIPEP
XPNPEP3

B

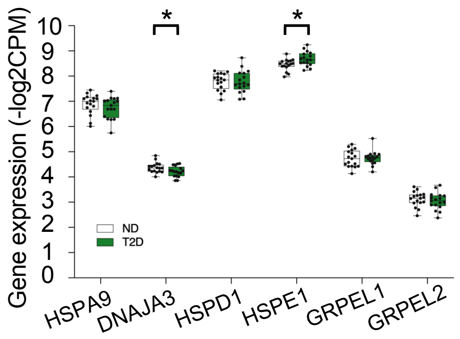
Gene expression (-log2CPM)
10
9
8
7
6
5
4
3
2
1
0
*
*
ND
T2D
HSPA9
DNAJA3
HSPD1
HSPE1
GRPEL1
GRPEL2

**Figure S4. Expression of mitochondrial proteases and chaperones is not altered in α-cells in human islet donors with T2D.**

Pseudobulk gene expression data, presented as log2CPM, of mitochondrial matrix proteases (A) and chaperones (B) from α-cells of human islet donors with or without T2D by single cell RNA sequencing. *$p < 0.05$ by both unpaired Student's two-tailed t-test and FDR < 5% for multiple testing correction. n = 17 non-diabetic donors, n = 17 donors with T2D.

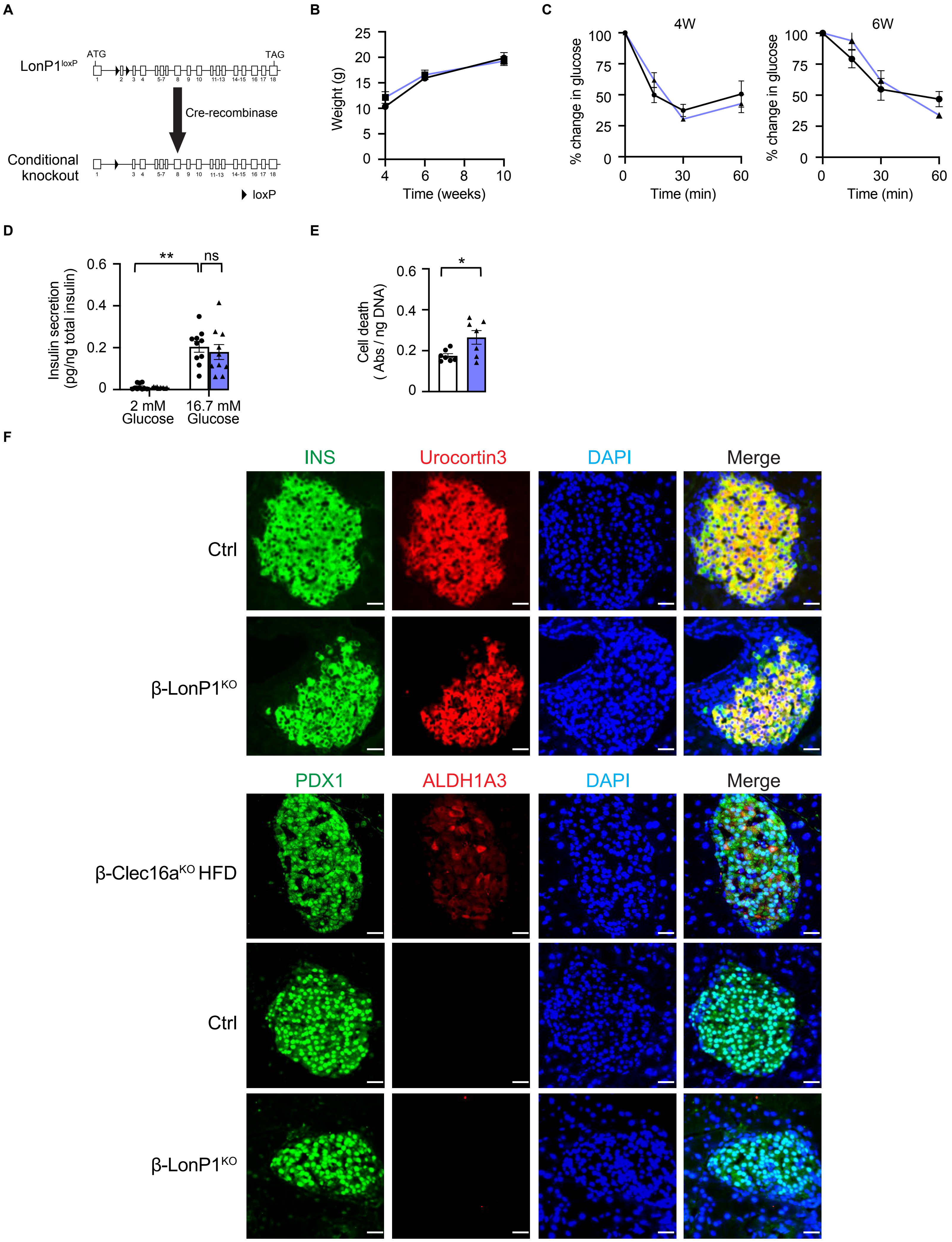
A
LonP1loxP
ATG
TAG
Cre-recombinase
Conditional knockout
loxP
B
Weight (g)
Time (weeks)
C
4W
6W
% change in glucose
Time (min)
D
Insulin secretion (pg/ng total insulin)
2 mM Glucose
16.7 mM Glucose
**
ns
E
Cell death ( Abs / ng DNA)
*
F
INS
Urocortin3
DAPI
Merge
Ctrl
β-LonP1KO
PDX1
ALDH1A3
DAPI
Merge
β-Clec16aKO HFD
Ctrl
β-LonP1KO

**Figure S5. LONP1 deficiency leads to reduced β-cell mass due to increases in β-cell apoptosis rather than alterations in β-cell maturity/dedifferentiation.**

(A) Model of Cre-mediated recombination of the *LonP1* locus. (B) Ad lib fed body weight measured in littermate control (Ctrl) and β-LonP1$^{KO}$ mice between ages 4-10 weeks. n = 5-8/group. (C) Blood glucose concentrations (presented as % of baseline glucose) measured during insulin tolerance testing (ITT) of 4-week-old (left) and 6-week-old (right) Ctrl and β-LonP1$^{KO}$ littermates. n = 6-9/group. (D) Glucose-stimulated insulin secretion following static incubations in 2 mM and 16.7 mM glucose, performed in isolated islets of 6-week-old Ctrl and β-LonP1$^{KO}$ littermates (normalized to total insulin content). n = 10 mice/group; $^{**}p < 0.01$ by one-way ANOVA followed by Tukey's multiple comparisons test. (E) Quantification of cell death by cytoplasmic histone-complexed DNA fragment ELISA (normalized to total DNA content) measured in isolated islets of 6-week-old Ctrl and β-LonP1$^{KO}$ mice. n = 7 mice/group; $^{*}p < 0.05$ by two-tailed Student's *t* test. (F) Representative immunofluorescence images (n = 5/group) depicting β-cell maturity (upper images) or dedifferentiation markers (lower images) from pancreatic sections of 6-week-old Ctrl and β-LonP1$^{KO}$ littermates, stained for insulin (green), Urocortin 3 (red) and DAPI (DNA - blue) or PDX1 (green), ALDH1A3 (red) and DAPI (DNA - blue). A representative image of pancreatic sections of high fat diet-fed of β-Clec16a$^{KO}$ mice, which were stained and imaged in parallel with Ctrl and β-LonP1$^{KO}$ sections, is shown as a positive control for ALDH1A3 immunostaining. Scale bar, 50 μm. n = 3-6 mice/group. All data in figure are presented as mean ± SEM.

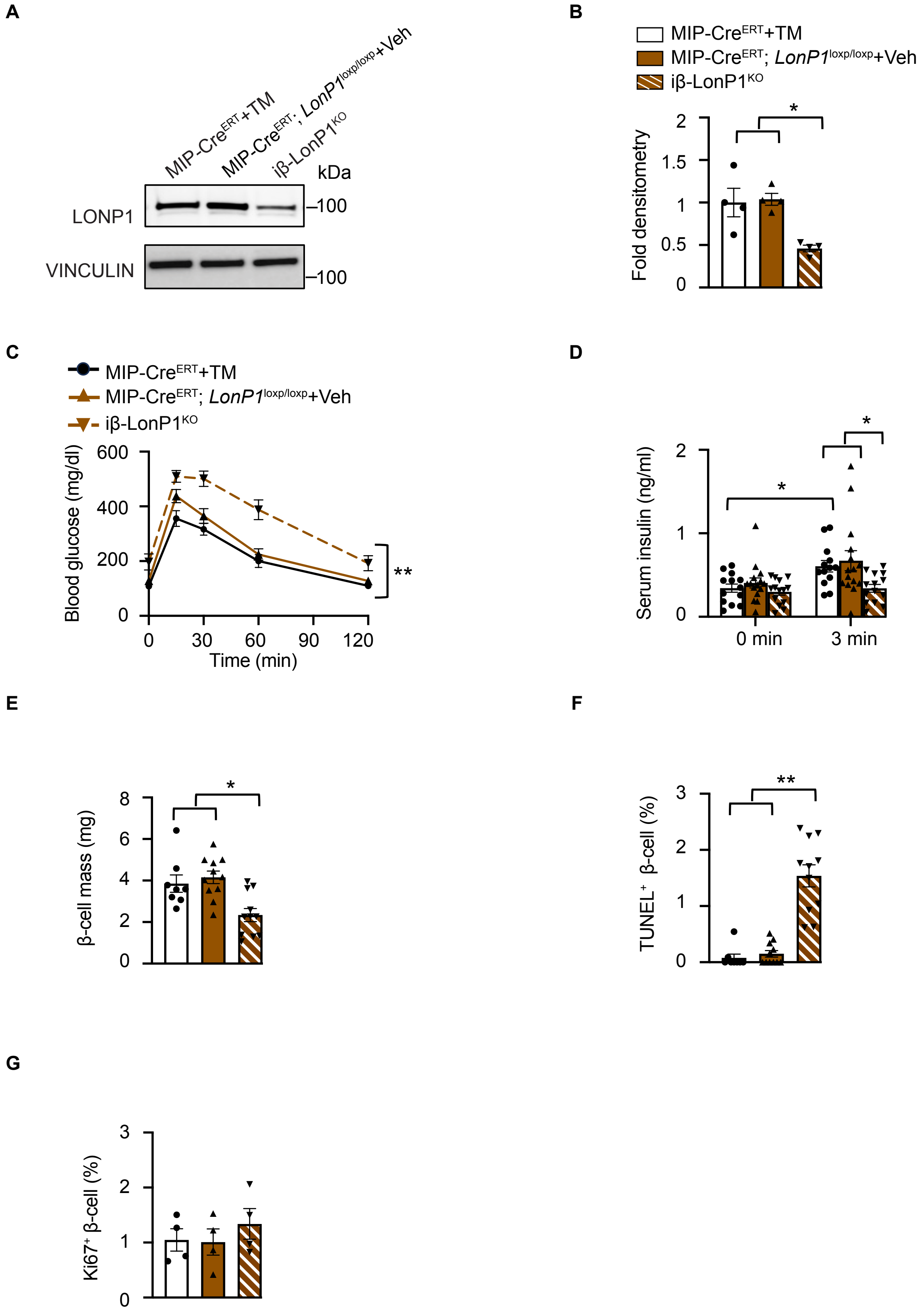
A
MIP-Cre^ERT+TM
MIP-Cre^ERT; LonP1^loxp/loxp+Veh
iβ-LonP1^KO
kDa
LONP1
–100
VINCULIN
–100
B
MIP-Cre^ERT+TM
MIP-Cre^ERT; LonP1^loxp/loxp+Veh
iβ-LonP1^KO
Fold densitometry
0
0.5
1
1.5
2
*
C
MIP-Cre^ERT+TM
MIP-Cre^ERT; LonP1^loxp/loxp+Veh
iβ-LonP1^KO
Blood glucose (mg/dl)
0
200
400
600
Time (min)
0
30
60
90
120
**
D
Serum insulin (ng/ml)
0
1
2
0 min
3 min
*
*
E
β-cell mass (mg)
0
2
4
6
8
*
F
TUNEL+ β-cell (%)
0
1
2
3
**
G
Ki67+ β-cell (%)
0
1
2
3

**Figure S6. Reductions in β-cell mass and glucose tolerance in adult mice following LONP1 deficiency are not due to developmental defects.**

(A) Expression of LONP1 evaluated by WB in islets isolated from 15-week-old iβ-LonP1$^{KO}$ mice as well as *MIP1*-Cre$^{ERT}$ + tamoxifen (TM) and MIP-Cre$^{ERT}$; *LonP1*$^{loxP/loxP}$ + Vehicle (Veh) control mice 7 weeks after Veh or TM administration. Representative images (left) of 4 mice/group with densitometry (normalized to VINCULIN) presented on graph (right). n = 4 mice/group; $^{*}p < 0.05$ by one-way ANOVA followed by Tukey's multiple comparisons test. (B) Blood glucose concentrations measured during IPGTT in 15-week-old iβ-LonP1$^{KO}$ mice as well as *MIP1*-Cre$^{ERT}$ + TM and MIP-Cre$^{ERT}$; *LonP1*$^{loxP/loxP}$ + Veh littermate controls 7 weeks after Veh or TM administration. n = 11-13 mice/group; $^{**}p < 0.01$ by one-way ANOVA followed by Tukey's multiple comparisons test. (C) Serum insulin measured during *in vivo* glucose-stimulated insulin release testing in 15-week-old iβ-LonP1$^{KO}$ mice as well as *MIP1*-Cre$^{ERT}$ + TM and MIP-Cre$^{ERT}$; *LonP1*$^{loxP/loxP}$ + Veh littermate controls 7 weeks after Veh or TM administration. n = 8-14 mice/group; $^{*}p < 0.05$ by one-way ANOVA followed by Tukey's multiple comparisons test. (D) β-cell mass measured in 15-week-old iβ-LonP1$^{KO}$ mice as well as *MIP1*-Cre$^{ERT}$ + TM and MIP-Cre$^{ERT}$; *LonP1*$^{loxP/loxP}$ + Veh littermate controls 7 weeks after Veh or TM administration. n = 8-11 mice/group; $^{*}p < 0.05$ by one-way ANOVA followed by Tukey's multiple comparisons test. (E) Quantification of β-cell apoptosis measured as the % of TUNEL$^{+}$/Insulin$^{+}$ cells performed in pancreatic sections of 15-week-old iβ-LonP1$^{KO}$ mice as well as *MIP1*-Cre$^{ERT}$ + TM and MIP-Cre$^{ERT}$; *LonP1*$^{loxP/loxP}$ + Veh littermate controls 7 weeks after Veh or TM administration. n= 8-11 mice/group; $^{**}p < 0.01$ by one-way ANOVA followed by Tukey's multiple comparisons test. (G) Quantification of β-cell replication measured as the % of Ki67$^{+}$/Insulin$^{+}$ cells performed in pancreatic sections of 15-week-old iβ-LonP1$^{KO}$ mice as well as *MIP1*-Cre$^{ERT}$ + TM and MIP-Cre$^{ERT}$; *LonP1*$^{loxP/loxP}$ + Veh littermate controls 7 weeks after Veh or TM administration. n = 4 mice/group. All data in figure are presented as mean ± SEM.

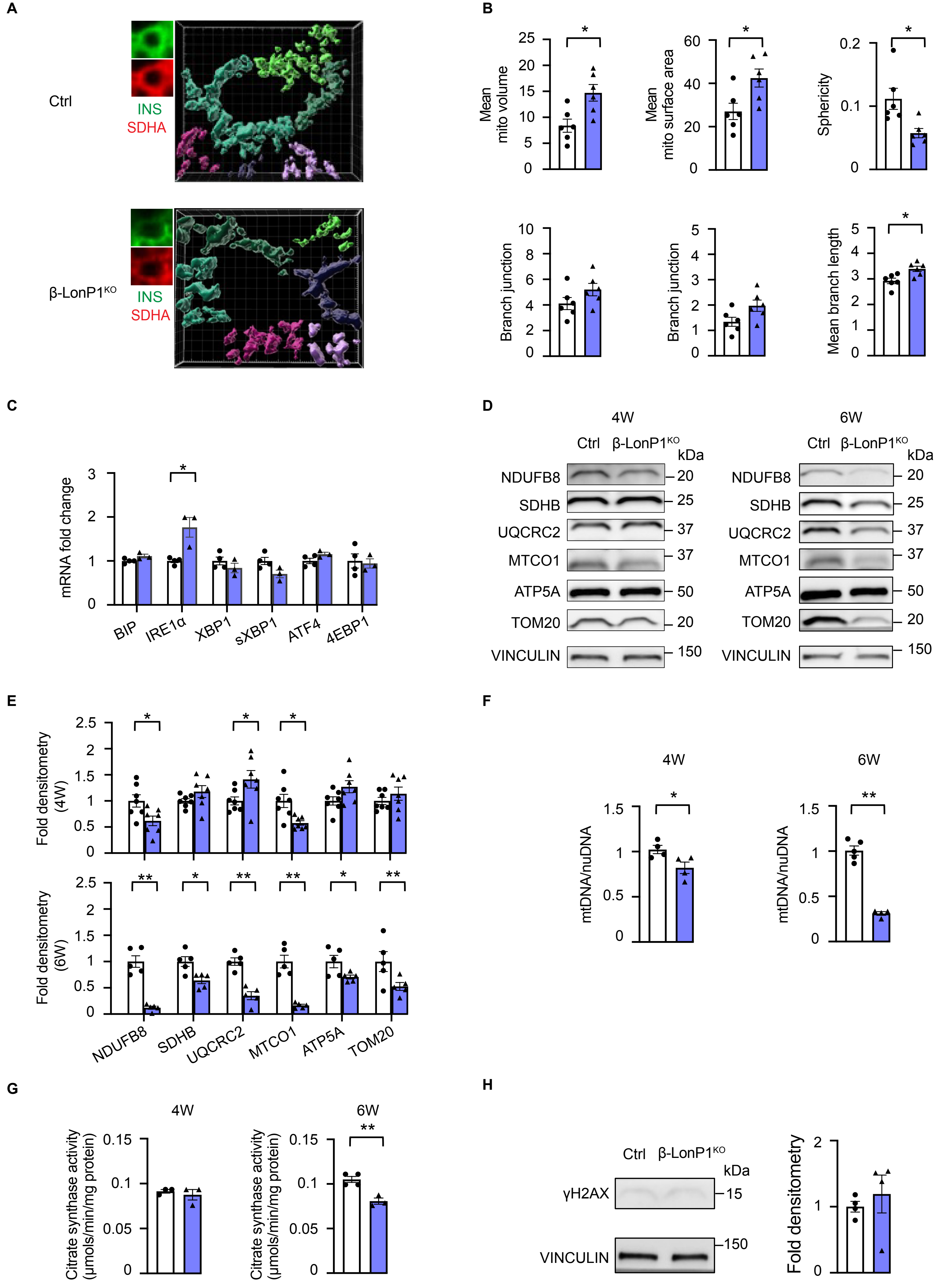

A
Ctrl
INS
SDHA
β-LonP1KO
B
Mean mito volume
Mean mito surface area
Sphericity
Branch junction
Mean branch length
C
mRNA fold change
BIP
IRE1α
XBP1
sXBP1
ATF4
4EBP1
D
4W
6W
Ctrl
β-LonP1KO
kDa
NDUFB8
SDHB
UQCRC2
MTCO1
ATP5A
TOM20
VINCULIN
E
Fold densitometry (4W)
Fold densitometry (6W)
NDUFB8
SDHB
UQCRC2
MTCO1
ATP5A
TOM20
F
4W
6W
mtDNA/nuDNA
G
4W
6W
Citrate synthase activity (µmols/min/mg protein)
H
Ctrl
β-LonP1KO
kDa
γH2AX
VINCULIN
Fold densitometry

**Figure S7. LONP1 deficiency leads to abnormal mitochondrial morphology and a decline of mitochondrial mass.**

(A) Imaris® generated three-dimensional reconstruction of deconvolution immunofluorescence Z-stack images at 60x magnification stained for insulin (green) and SDHA (red) in pancreatic sections of 6-week-old Ctrl and β-LonP1$^{KO}$ mice. Representative images of 7 independent mice/group. Each unique color represents a separate β-cell mitochondrial network cluster. (B) β-cell mitochondrial morphology and network analysis of deconvolution immunofluorescence Z-stack images at 100X magnification stained for SDHA (and insulin) from pancreatic sections of 6-week-old Ctrl and β-LonP1$^{KO}$ mice by Mitochondria Analyzer. n = 6 mice/group. $^{*}p < 0.05$ by two-tailed Student's *t* test. (C) Quantitative RT-PCR of markers of ER stress, normalized to *HPRT* expression, from RNA isolated from 6-week-old Ctrl and β-LonP1$^{KO}$ islets. n = 3-4/group. $^{*}p < 0.05$, $^{**}p < 0.01$ by two-tailed Student's *t* test. (D) Expression of OXPHOS complex subunits and TOM20 by WB in isolated islets from Ctrl and β-LonP1$^{KO}$ mice at both 4-weeks (left) and 6-weeks (right) of age. Representative images of 5-7 mice/group. VINCULIN serves as a loading control. (E) Quantification of OXPHOS complex subunits and TOM20 levels in Ctrl and β-LonP1$^{KO}$ islets at both 4-weeks (upper) and 6-weeks (lower) of age by densitometry (normalized to VINCULIN) from studies in Figure S7D. n = 5-7 mice/group. $^{*}p < 0.05$, $^{**}p < 0.01$ by two-tailed Student's *t* test. (F) Relative mtDNA content normalized to nuclear DNA expression measured by qPCR in isolated islets of Ctrl and β-LonP1$^{KO}$ littermates at both 4-weeks (left) and 6-weeks (right) of age. n = 4-5 mice/group. $^{*}p < 0.05$, $^{**}p < 0.01$ by two-tailed Student's *t* test. (G) Citrate synthase activity measured in isolated islets of Ctrl and β-LonP1$^{KO}$ littermates at both 4-weeks (left) and 6-weeks (right) of age. n = 3-4 mice/group. $^{**}p < 0.01$ by two-tailed Student's *t* test. (H) Expression of phospho-γH2AX levels by WB in isolated islets of 4-week-old Ctrl and β-LonP1$^{ko}$ littermates. Representative images (left) of 4 mice/group. Quantification of phospho-γH2AX levels (normalized to VINCULIN) shown in graph on right. n = 4 mice/group. All data in figure are presented mean ± SEM.

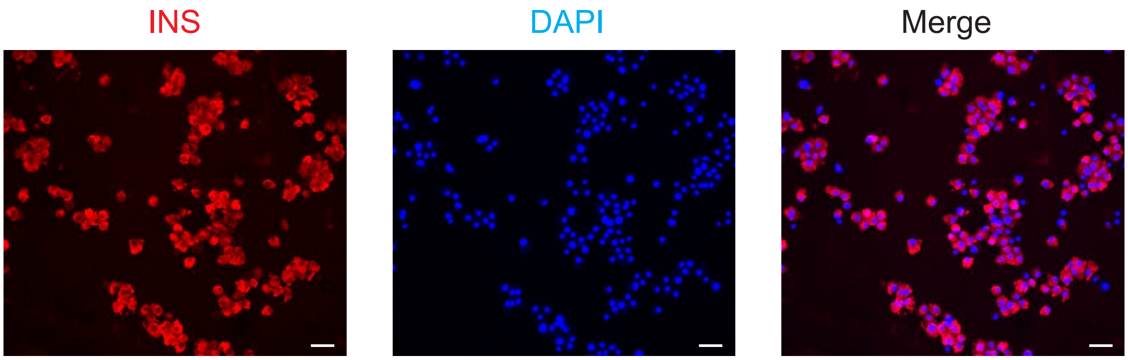
INS
DAPI
Merge

**Figure S8. Generation of human β-cell enriched pseudoislets.**

Immunofluorescence imaging performed in human β-cell enriched pseudoislets, generated by magnetic sorting for the β-cell surface marker NTPDase3, following dissociation for cytocentrifugation and imaging, stained for insulin (red) and DAPI (DNA - blue). Scale bar, 50 μm. Representative image of 4 β-cell enriched pseudoislet preparations each from independent human islet donors.
